# Functional and Structural Basis of Omicron BA.3.2.1 Spike

**DOI:** 10.64898/2026.02.02.702852

**Authors:** Yan Wang, Yanping Hu, Zhenhang Chen, Jing Zou, Ke Zhang, Ping Ren, Pei-Yong Shi, Bo Liang, Xuping Xie

**Author notes:** Y.W. and Y.H. contributed equally to this work. Corresponding authors: X.X.

## Abstract

SARS-CoV-2 Omicron descendant BA.3.2, derived from BA.3 and carrying 39 additional spike substitutions, prompts concerns about altered fitness and antigenicity. Using BA.3.2.1 as a representative sublineage, we have engineered live-attenuated mNeonGreen SARS-CoV-2 encoding spikes from BA.3.2.1 and the JN.1-derived LP.8.1 and XEC, and benchmarked them against BA.3 and KP.3 in Calu-3 cells and primary human airway epithelium. BA.3.2.1 outcompetes BA.3 yet replicates more slowly than JN.1-descendants. Despite increased RBD-hACE2 affinity, BA.3.2.1 spike binds hACE2 less efficiently than LP.8.1, and shows the greatest resistance to neutralization by sera from KP.2/KP.3 infections. Cryo-EM reveals a predominantly closed, compact, asymmetric trimeric spike with a remodeled N-terminal domain, a distinct fusion-peptide-proximal region and altered N-glycosylation. BA.3.2.1 spike adopts a fine-tuned receptor-binding interface and shifted monoclonal-antibody footprints. These findings identify a trade-off between immune evasion and replication fitness driven by closed spike conformation, rationalizing BA.3.2’s limited prevalence and underscoring the necessity for continued variant surveillance.

## Introduction

Severe acute respiratory syndrome coronavirus 2 (SARS-CoV-2), the etiologic agent of coronavirus disease 2019 (COVID-19) pandemic, has resulted in unprecedented global morbidity and mortality since its emergence in December 2019^1,2^. Continued antigenic drift and episodic saltations have produced successive variant waves with enhanced immune evasion and altered transmissibility, most notably after the emergence of Omicron and its sublineages^3^. The initial Omicron BA.1 marked a major evolutionary shift characterized by extensive spike mutations and broad immune escape^4–7^. Subsequent sublineages (BA.2, BA.3, BA.4, BA.5) and Omicron-associated recombinants such as XBB drove additional global waves of infection^8–11^. In mid-2023, the highly divergent BA.2.86 lineage appeared, with JN.1 subsequently becoming globally predominant and giving rise to multiple descendants, including KP.2/KP.3, and later LP.8.1, NB.1.8.1, and XFG. The JN.1 descendants shaped SARS-COV-2 circulation through 2024-2025^12–16^.

In parallel, BA.3.2 emerged as a long-branch descendant of the rarely circulating BA.3. BA.3.2 was first detected in South Africa (November 2024) and designated by WHO as a Variant Under Monitoring in Dec 2025^17^. Since its emergence, BA.3.2 has drawn substantial attention for an unusually high burden of spike substitutions (>39 additional spike mutations versus BA.3 and >75 changes distinct from contemporaneous JN.1 descendants such as LP.8.1 and XFG)^12,18^. Although prevalence remains low, BA.3.2 has been reported across Africa, Europe, the United States, and Oceania^19^, and continues to spread to many other countries, where KP.3, LP.8.1, or XFG circulate. Understanding how BA.3.2 mutations reshape spike structure and function is therefore critical for predicting viral fitness and immune escape.

SARS-CoV-2 is an enveloped, positive-sense single-stranded RNA virus whose surface is studded with trimeric spike (S) glycoproteins. The spike mediates viral entry by binding to the host cell receptor angiotensin-converting enzyme 2 (ACE2)^20,21^. The spike comprises two subunits, S1 and S2, separated by a furin cleavage site. The S1 subunit contains the N-terminal domain (NTD), receptor-binding domain (RBD), and subdomains SD1 and SD2. RBD often toggles between “down” (ACE2-inaccessible) and “up” (ACE2-accessible) conformations^22–24^. Distal regions in the S protein, including SD1 and SD2, allosterically influence RBD positioning. The S2 subunit includes the S2′ site (typically cleaved by TMPRSS2 or other host serine protease), fusion peptide (FP), fusion-peptide proximal region (FPPR), heptad repeats (HR1 and HR2), central helix (CH), connector domain (CD), transmembrane domain (TM), and cytoplasmic tail (CT)^25^. ACE2 binding and cleavages at the furin and S2’ sites trigger major conformational rearrangements that drive membrane fusion^20^. Conformational “gating” of the RBD-up state, together with S1/S2 processing, can therefore modulate hACE2 engagement.

Spike is the primary target of vaccines and therapeutic neutralizing antibodies (NAbs). Most potent NAbs often target RBD. Following the Barnes framework^26^, RBD-directed NAbs are commonly grouped into distinct classes: Class I overlaps the ACE2-binding site and engages only RBD-up^27^; Class II also compete with ACE2 but can bind RBD-up and RBD-down^26^; Class III targets the outer, ACE2-distal face and often more conserved surfaces^28–30^; and Class IV recognizes cryptic epitopes exposed as the RBD opens^31,32^. Class V maps to additional cryptic epitopes opposite the ACE2 interface^33^. Moreover, NTD- and S2-directed NAbs provide complementary neutralization mechanisms^34–37^.

Prior work suggests that BA.3.2 exhibits pronounced immune evasion yet reduced infectivity, fusogenicity, and replication capacity compared with JN.1-descendant variants^12,18,19,38^. However, many studies rely on pseudovirus systems with limited analyses using authentic virus in physiologically relevant primary human airway epithelium (HAE). A rigorous functional-structural definition of BA.3.2, benchmarked against contemporaneous JN.1 descendants, is therefore needed to understand how its extensive spike mutations reshape receptor engagement, conformational dynamics, and antigenicity, and contextualize its persistent but low observed prevalence.

In this study, we investigated the functional role and structural features of BA.3.2.1 as a representative sublineage. We engineered attenuated mNeonGreen (mNG) reporter SARS-CoV-2 encoding spikes from BA.3.2.1 or co-circulating JN.1-derived strains (LP.8.1, XEC), and benchmarked them against BA.3 and KP.3 in Calu-3 cells and primary HAE cultures. We profiled neutralization using convalescent sera from KP.2/KP.3 infections. Finally, we determined cryo-EM structures of BA.3.2.1 and LP.8.1 spikes in apo and hACE2-bound states. Our findings reveal a trade-off between BA.3.2’s immune escape and replication fitness, providing a structural-functional rationale for BA.3.2’s limited global expansion.

## Results

### Generation and characterization of BA.3.2.1-spike mNG SARS-CoV-2

BA.3.2 emerged in late 2024, while JN.1-descendant variants co-circulated and followed a successive epidemiological pattern: KP.3→XEC→LP.8.1. At the time this study was completed, XFG and NB.1.8.1 had become the dominant strains. Two BA.3.2 sublineages, BA.3.2.1 (containing P681R and P1162R) and BA.3.2.2 (containing K356T, A575S, and P681H), were reported. This study focused on BA.3.2.1 due to its earlier emergence. **Fig. 1a** depicts the phylogenetic relationships among these variants. Defining the functional role and immune evasion capacity of the BA.3.2 spike, in comparison with JN.1-descendant spikes (KP.3, XEC, LP.8.1), could help explain why BA.3.2 did not outcompete other circulating strains. To address this, we engineered spike genes from BA.3.2.1, XEC, and LP.8.1 into the mNeonGreen (mNG) SARS-CoV-2 backbone (**Extended Data Fig. 1a-b**). This reporter virus, derived from strain USA-WA1/2020 (WA1) with open reading frame 7 (ORF7) replaced by mNG, is highly attenuated *in vivo* and suitable for safe use in BSL-3 for variant analysis^13,39,40^. All three variant-spike mNG SARS-CoV-2s were robustly recovered in Vero E6-TMPRSS2 cells, with P1 stock titers exceeding 10^6^ PFU/mL. For comparison, BA.3-spike and KP.3-spike mNG SARS-CoV-2 from previous studies were included^4,40^. All tested Omicron spike variants produced plaques of similar size, though noticeably smaller than those formed by WA1-spike mNG SARS-CoV-2 (**Extended Data Fig. 2a**).

**Fig. 1:**
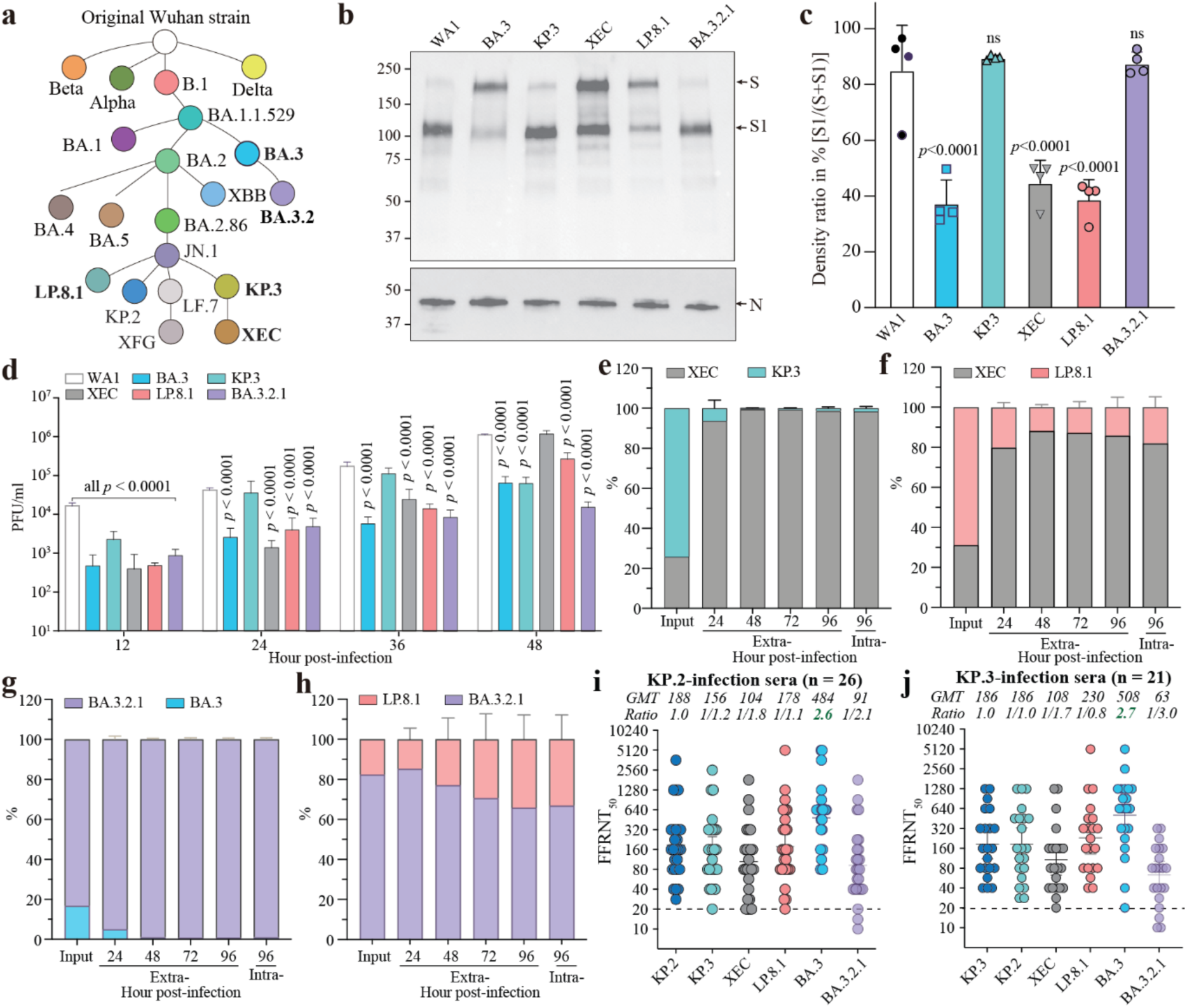
Characterization of mNG SARS-CoV-2 spike variants. **a,** Phylogenetic relationship of JN.1 and BA.3 descendants, adapted from CDC data (www.cdc.gov/covid/php/variants/ variants-and-genomic-surveillance.html). **b,** Western blot analysis of spike and nucleocapsid protein in virions. S, full-length spike; S1, N-terminal Furin cleavage fragment of spike; N, nucleocapsid. **c,** Ratio of S1 to total spike (S plus S1) in virions. Means and standard deviations from four independent experiments are shown. *P* values (one-way ANOVA with multiple comparison corrections) are shown. **d,** Growth kinetics of mNG SARS-CoV-2 spike variants on Calu-3 cells. Means and standard deviations from four independent experiments are shown. Each group was compared with WA1 using two-way ANOVA with Dunnett’s multiple comparison corrections. *P* values are indicated. **e-h,** Competition analysis of spike variants on HAE. The percentage of XEC- and KP.3-spike RNA (**e**), XEC- and LP.8.1-spike RNA (**f**), BA.3.2.1- and BA.3-spike RNA (**g**), LP.8.1- and BA.3.2.1-spike RNA (**h**) are shown. Means ± standard deviations from five independent experiments are presented. **i-j**, FFRNT_50_ of KP.2- (**i**) or KP.3-infection (**j**) sera against mNG SARS-CoV-2 spike variants. Solid lines and numeric values above each panel indicate GMTs. Error bars show 95% confidence intervals. GMT ratios are shown for each variant relative to KP.2-spike or KP.3-spike. Dotted lines indicate the FFPNT limit of detection. Group comparisons were performed using the Wilcoxon matched-pairs signed-rank test. The *P* values (determined using the Wilcoxon matched-pairs signed-rank test) for group comparison of GMTs are shown in Supplementary Table 3.

We first characterized spike S1/S2 cleavage across the variants. Virions released from Vero E6 cells infected with each spike variant mNG SARS-CoV-2 were analyzed by Western blot using anti-S1 and anti-N antibodies. Two distinct cleavage patterns were observed: KP.3 and BA.3.2.1 spikes were predominantly cleaved (>80%), comparable to the WA1 spike, whereas BA.3, XEC, and LP.8.1 spikes exhibited markedly reduced cleavage (∼40%) (**Fig. 1b-c**). Next, we examined replication kinetics of these spike variants in the human lung adenocarcinoma cell line Calu-3 (**Fig. 1d**). As expected, Omicron spike variants overall replicated more slowly than WA1 spike mNG SARS-CoV-2. Notably, JN.1 descendants displayed a replication trend of KP.3>XEC>LP.8.1, which contrasted with their epidemiological succession. BA.3- and BA.3.2.1-spike mNG SARS-CoV-2 were generally more attenuated than JN.1 descendants, with BA.3.2.1-spike showing the greatest attenuation.

### BA.3.2.1 spike is more attenuated than JN.1 descendants in HAE

To compare replication fitness under physiologically relevant conditions, we conducted paired competition assays in primary human airway epithelial (HAE) cultures. Equal PFUs of two spike-variants were mixed and used to infect HAE cells, and variant proportions were quantified by next-generation sequencing (NGS) from daily apical washes and endpoint intracellular RNA. Four variant pairs were selected based on evolutionary relationships and epidemiological relevance: KP.3 vs. XEC, XEC vs. LP.8.1, BA.3 vs. BA.3.2.1, and LP.8.1 vs. BA.3.2.1. Although PFU inputs were normalized, initial RNA ratios varied (**Extended Data Fig. 2b-e**), indicating infectivity differences.

Competition outcomes revealed marked differences in replication fitness among spike variants in HAE. Among JN.1 descendants, XEC-spike rapidly outcompeted KP.3-spike, increasing from 25% at input to >90% by 24-96 h post-infection (p.i.) (**Fig. 1e, Extended Data Fig. 2b**). XEC-spike also dominated LP.8.1-spike, rising from 30% to >80% over 24-96 h p.i. (**Fig. 1f, Extended Data Fig. 2c**). BA.3.2.1-spike strongly outgrew BA.3-spike (**Fig. 1g, Extended Data Fig. 2d**). However, in competition with LP.8.1-spike, BA.3.2.1-spike showed reduced replication, with its relative abundance declining from 24-96 h p.i. (**Fig. 1h, Extended Data Fig. 2e**).

### BA.3.2.1-spike exhibits the greatest immune escape among tested variants

Next, we evaluated the sensitivity of these spike variants to human convalescent sera. Serum samples were collected from 47 adult participants at UTMB clinics, including 26 individuals with documented KP.2 infections 21-99 days (median 42.5 days) before sampling (KP.2-infection panel in **Supplementary Table 1**) and 21 individuals with KP.3 infections 23-105 days prior (median 50 days) (KP.3-infection panel in **Supplementary Table 2**). Neutralizing activity was assessed using a fluorescent focus reduction neutralization test (FFRNT), in which each serum sample was tested in parallel against these spike variants plus the KP.2-spike mNG SARS-CoV-2^40^.

KP.2-infection panel sera neutralized KP.2-, KP.3-, XEC-, LP.8.1-, BA.3-, BA.3.2.1-spike mNG SARS-CoV-2 with geometric mean titers (GMT) of 188, 156, 104, 178, 484, and 91 (**Fig. 1i**), respectively. Similarly, KP.3-infection panel sera showed GMTs of 186, 186, 108, 230, 508, and 63 against the same variants (**Fig. 1j**). KP.2- and KP.3-spike exhibited similar sensitivity to both panels, consistent with previous findings^40^. BA.3-spike was most sensitive (2.6-2.7× higher GMTs than KP.2-/KP.3-spike), whereas BA.3.2.1-spike showed the strongest resistance (2.1-3.0× lower GMTs). XEC-spike displayed modest resistance (1.7-1.8× lower GMTs), while LP.8.1-spike remained comparable to KP.2-/KP.3-spike. These results indicate BA.3.2.1-spike has the greatest escape from neutralization elicited by prior KP.2 and KP.3 infection.

### BA.3.2.1 spike exhibits less structural flexibility than LP.8.1 spike

To define the structural basis for BA.3.2.1 spike-mediated fitness and immune evasion, we determined its trimeric structure by single-particle cryo-EM (**Extended Data Fig. 3**). Most BA.3.2.1 spike particles (82.5%) adopted a closed 3-RBD-down conformation (3.2-3D; PDB 9YSJ, EMD 73393), while a minor fraction displayed a flexible state (3.2-1M; PDB 9YX6, EMD 73395) with two down RBD and one unresolved RBD (**Fig. 2a**). These structures were resolved at 2.6 Å and 3.0 Å, respectively (**Extended Data Fig. 3a-e, Supplementary Table 5**). For comparison, we also determined the cryo-EM structure of the LP.8.1 spike (**Extended Data Fig. 4a-f, Supplementary Table 6**). Similar to other JN.1 descendants (e.g., JN.1.11 and KP.3.1.1)^41^, LP.8.1 spike particles displayed three conformational states: closed (3-RBD-down, 36.3%; 8.1-3D; PDB 9YSK, EMD 73394), open (1-RBD-up, 35.8%; 8.1-1U; PDB 9YW0, EMD 73535), and flexible (2-RBD-unresolved, 27.9%; 8.1-2M; EMD 73396) with resolutions of 2.5-2.6 Å (**Fig. 2b)**.

**Fig. 2:**
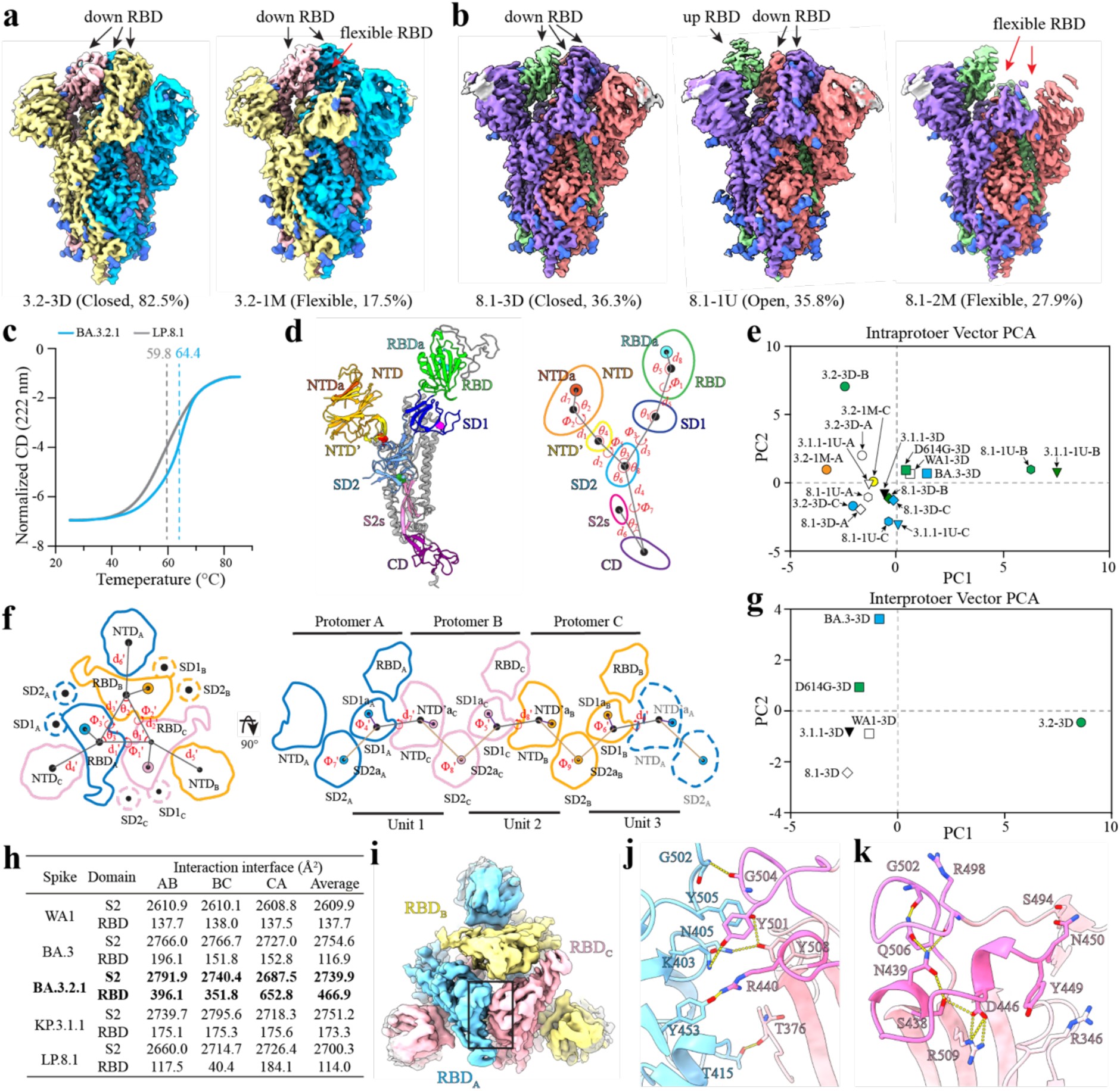
Structures of BA.3.2.1 and LP.8.1 spikes. **a,** Cryo-EM maps of BA.3.2.1 spike in closed state (left) and flexible state (right). Protomers are shown in light blue, yellow, and light pink, respectively. **b,** Cryo-EM maps of LP.8.1 spike in closed state (left), open state (middle), and flexible state (right). Protomers are shown in light magenta, green, and salmon, respectively. **c,** Melting curves of BA.3.2.1 and LP.8.1 spikes determined by CD analysis. **d**, Intraprotomer vector analysis. Left: atomic model of protomer A from closed BA.3.2.1 spike highlighting vector positions. Right: Schematic annotating vectors, angles, and dihedrals between different structural elements. **e,** PCA of the intraprotomer vector magnitudes, angles, and dihedrals. **f**, Interprotomer vector analysis. Left, top-down view of the spike protein highlighting the positions of S1 subunit domains. Right, a two-dimensional representation of the domains showing the interprotomer vectors, angles, and dihedrals between them. **g,** PCA of the interprotomer vector magnitudes, angles, and dihedrals. **h**, The binding surface areas between neighboring S2 subunits and the RBDs in spikes with closed conformations. **i**, Top view of the closed BA.3.2.1 spike cryo-EM map, highlighting the inter-RBD interaction between RBD_A_ and RBD_C_ (black box). **j**, Interaction networks between RBD_A_ and RBD_C_. The 497-505 and 438-451 loops are colored plum and violet, respectively. Dashed lines represent hydrogen bonds. **k**, Interaction with RBD_C_ that stabilized the remodeled 497-505 loop and the 438-451 loop. Dashed lines represent hydrogen bonds or salt bridges.

To further assess structural heterogeneity and conformational dynamics, we performed 3D variability analysis (3DVA) of both cryo-EM datasets. BA.3.2.1 spike in the 3-RBD-down state exhibited limited conformational heterogeneity, showing localized low-amplitude variability primarily at the HR2 region with minor excursions in the NTDs and RBDs (**Extended Data Fig. 5a-b; Supplementary Movie 1**). The cryo-EM structure of the flexible BA.3.2.1 spike overall resembled the closed conformation (RMSD = 0.395 Å) (**Extended Data Fig. 5c**), while one RBD was unresolved, consistent with higher-amplitude variability and more conformational heterogeneity revealed by 3DVA (**Extended Data Fig. 5d-e; Supplementary Movie 2**). In contrast, when comparing 3-RBD-down particles, LP.8.1 spikes displayed greater conformational heterogeneity than BA.3.2.1, with more pronounced amplitude variability across the RBDs and NTDs (**Extended Data Fig. 5f-g, Supplementary Movie 3**), indicating enhanced breathing. The open and flexible LP.8.1 spike particles showed a similar pattern of variability to the closed ones, indicating an overall increase in conformational variability in LP.8.1 (**Extended Data Fig. 5h-k, Supplementary Movie 4-5**). Consistent with these observations, circular dichroism (CD) analysis revealed that BA.3.2.1 spikes were more thermostable, with a melting temperature 4.6°C higher than LP.8.1 (**Fig. 2c**). Together, these findings demonstrate that the prefusion BA.3.2.1 spike exhibited reduced conformational heterogeneity compared with LP.8.1.

### Quaternary structure of BA.3.2.1 spike

Alignment of individual protomers revealed pronounced structural variations in SD1 and the RBD among BA.3.2.1 spike protomers, in contrast to the minimal variability observed for LP.8.1 (**Extended Data Fig. 5l-m**). To quantitatively compare their structural differences, we applied a vector-based quaternary analysis as described previously^42,43^. Briefly, we assigned a central coordinate to each spike domain and computed intra-protomer descriptors, angles, dihedrals, and distances between domain vectors (**Fig. 2d**). The comparative set included spike structures from WA1 (PDB 6VXX), BA.3 (PDB 7XIY), D614G (PDB 7K4E), and KP.3.1.1 (PDB 9ELI, closed; 9ELH, open). For cross-structure comparison, the protomer with the greatest conformational deviation was designated protomer B; A and C were then labeled clockwise from B when viewed from the extracellular side.

Principal component analysis (PCA) was performed using the 25 intra-protomer geometric descriptors (10 distances, 8 angles, 7 dihedrals) across variants (**Supplementary Table 7**). PC1 reported global hinge/torsional openness with the largest loadings from Φ_3_/Φ_1_ (+) and θ_6_/θ_5_/θ_1_ (-), and modest contribution by d_5_ and Φ_2_ (+); whereas PC2 indexed orthogonal spacing/auxiliary changes, dominated by d_3_/d_10_ (+) and opposed by Φ_5_ (-), with θ_4_/θ_2_ (+), and θ_8_ (-) also contributing. This PC1-PC2 map resolved two state-driven clusters: a left-hand RBD down-state cluster and a right-hand RBD up-state cluster (**Fig. 2e**). Within this space, BA.3.2.1-closed (3.2-3D) localized to a compact hinge and a distinct SD1-SD2 spacing/auxiliary configuration (centroid: PC1, −2.04; PC2, 2.45), and showed marked protomer asymmetry along PC2 (A: −1.63, 2.00; B, −2.44, 7.05; C, −2.06, −1.70), with protomer B showing the largest spacing/auxiliary shift (**Fig. 2e**). By contrast, LP.8.1-Open (1.51, −1.00) and KP.3.1.1-Open (2.10, −0.84) populated the RBD-up cluster, with their open-RBD protomers (8.1-1U-B and 3.1.1-1U-B) lying at the positive extreme of PC1, consistent with increased hinge openness. LP.8.1-closed (8.1-3D) remained compact with a different auxiliary configuration. BA.3.2.1-Flexible (3.2-1M) overall resembled the closed geometry for the two resolved protomers. WA1 (WA1-3D), D614G (D614G-3D), and BA.3 (BA.3-3D) lay near the origin, with modest hinge opening and mild spacing changes.

Consistently, descriptor-level inspection (**Extended Data Fig. 6a-c**) revealed measurable differences among the three protomers in closed BA.3.2.1. Protomer C showed θ_1_ (SD2-SD1-RBD) and θ_5_ (SD1-RBD-RBDa) shift opposite the open-state ensemble, indicating a more down-oriented RBD, accompanied by lower Φ_1_ (SD2-SD1-RBD-RBDa) and Φ_3_ (NTD’-SD2-SD1-RBD), consistent with a more twisted RBD-SD2/NTD’ relationship. Protomer B exhibited outward SD1 rotation (reduced θ_3_ [NTD’-SD2-SD1] and θ_8_ [CD-SD2-SD1]), with corresponding Φ_5_/Φ_7_/d_3_ shifts (greater SD1 mobility), and a modest RBD elevation relative to A/C (lower θ_1_, increased d_5_, and decreased d_9_). These placements supported a compact intra-protomer hinge in BA.3.2.1-closed with a distinct SD1-SD2 auxiliary configuration.

To assess subdomain organization across protomers at the closed state, we performed inter-protomer vector analysis (**Fig. 2f**) by defining SD1/RBD-NTD/NTD’ conformational units and measuring their trimer-spanning disposition with SD2 as previously described^42,43^. We included previously reported closed spikes of WA1 (PDB 6VXX), BA.3 (PDB 7XIY), D614G (PDB 7K4E), and KP.3.1.1 (PDB 9ELI) for this analysis. PCA of cross-protomer descriptors (d_1_’-d_9_’, θ_1_’-θ_3_’, Φ_1_’-Φ_9_’) resolved variant-specific trimer arrangements (**Fig. 2g, Supplementary Table 8**). PC1 (73.0%) predominantly captured inter-RBD compaction/expansion, with the largest loadings from d_1_’-d_3_’ and RBD-centric torsions/angles (Φ_1_’-Φ_3_’/θ_1_’-θ_3_’); whereas PC2 (16.6%) indexed orthogonal SD1/SD2-NTD’ coupling (dominated by Φ_5_’-Φ_7_’ and d_7_’-d_9_’). In this space, BA.3.2.1-closed occupied the PC1-positive extreme, indicating a quaternary compaction outlier with contracted inter-RBD spacing and constrained RBD-centric torsions. By contrast, BA.3 lay at the PC2-positive extreme, consistent with pronounced SD1/SD2-NTD’ rearrangements. LP.8.1 and KP.3.1.1 were both PC1- and PC2-negative, reflecting expanded inter-protomer RBD spacing with comparatively reduced orthogonal torsion, whereas D614G and WA1 occupied intermediate positions.

Descriptor-level inter-protomer analysis (**Extended Data Fig. 6**) corroborated these axes. BA.3.2.1 showed a contracted inter-RBD triad (reduced d_1_’-d_3_’, Φ_1_’ and increased Φ_2_’/Φ_9_’), and angle shifts consistent with a tightened RBD arrangement (decreased θ_3_’/θ_1_’ and increased θ_2_’). BA.3.2 exhibited pronounced cross-protomer SD1-NTD’ rearrangements (increased d_7_’-d_9_’/Φ_5_’-Φ_7_’); whereas LP.8.1 and KP.3.1.1 displayed expanded inter-RBD spacing (d_1_’-d_3_’) with reduced SD1/SD2-NTD’ coupling (reduced Φ_5_’-Φ_7_’/d_7_’-d_9_’). D614G and WA1 were intermediate. Together, the intra- and inter-protomer maps converged on BA.3.2.1-closed as a compact, “locked” quaternary architecture with contracted inter-RBD geometry, compact intra-protomer hinging and a distinct SD1/SD2 auxiliary signature.

### Unique RBD-RBD interface in closed BA.3.2.1 spike

The compacted structures of BA.3.2.1 spike prompted us to examine the RBD-RBD interfaces closely. PISA analysis revealed that BA.3.2.1 spike harbored a substantially larger RBD-RBD interface than other variants’ spikes, including LP.8.1, while S2 contact areas remained similar (**Fig. 2h**). The most extensive interface occurred between RBDs A and C, which was supported by the cryo-EM maps (**Fig. 2i**). Local refinement identified multiple induced inter-RBD interactions, including T376 (RBD_C_)-T415 (RBD_A_), R440 (RBD_C_)-Y453 (RBD_A_), Y501 (RBD_C_)-K403 (RBD_A_), G504 (RBD_C_)-G502 (RBD_A_), and Y508 (RBD_C_)-N405/Y505 (RBD_A_) (**Fig. 2j**). Accompanying those contacts, RBD_C_ underwent pronounced remodeling: the 497-505 loop bent toward the interface; the 438-451 segment reorganized into two shorter helices (residues 438-440 and 446-448) flanking a protruding loop (residues 441-445) extending toward RBD_A_ (**Extended Data Fig. 7a**). Additional intramolecular networks formed within this loop and adjacent residues (**Fig. 2k**), further stabilizing the conformational configuration. The RBD_B_ 443-450 loop was also slightly re-orientated (**Extended Data Fig. 7a**). However, such structural remodeling was absent in LP.8.1, whose RBDs maintained highly similarity across protomers and closely matched WA1 and KP.3 RBDs (pairwise RMSD 0.617-0.692 Å), aside from minor, Omicron-conserved variations in the 365-373 loop due to convergent mutations S371F, S373P, and S375F (**Extended Data Fig. 7b**)^44,45^. Collectively, these features explain enhanced RBD packing, restricted RBD mobility, and clamping the BA.3.2.1 spike in a tightly closed conformation.

### BA.3.2.1 NTD remodeling

Beyond the RBD, BA.3.2.1 NTD exhibited pronounced structural remodeling. The NTD is another important target of NAbs, with most recognizing the NTD-1 antigenic supersite formed by five surface-exposed loops: N1 (residues 14-26), N2 (residues 67-79), N3 (residues 141-156), N4 (residues 177-186), and N5 (residues 246-260) (**Extended Data Fig. 8a**)^46^. To reconstruct the NTD-1 supersite in BA.3.2.1 and LP.8.1, we performed local refinement (**Extended Data Fig. 3i, 4g, Supplementary Table 9-10**). In BA.3.2.1 NTD (PBD 9Z3Y), N1, N2, and N5 remained unresolved in the local refinement map and were subsequently modeled using Phenix’s predict-and-build workflow guided by AlphaFold^47,48^. A similar approach was applied to LP.8.1 NTD (PBD 9YT7) to model unresolved N3, N4, and N5.

Structural superimposition revealed substantial differences between LP.8.1 and BA.3.2.1 NTDs (**Fig. 3a-b**). In LP.8.1, N1 and N2 were well resolved (**Extended Data Fig. 8a**). The 17-20 MPLF insertion, together with Δ24-27 and Δ69-70 deletions, positioned N1 to thread through N2, stabilizing a unique N1-N2 crossing via a C15-C131 disulfide bond, hydrogen bonds between N21 and N74/R78, and two hydrophobic interaction networks involving F79-P17-F139 and F18-L244 (**Fig. 3c, Extended Data Fig. 8b**). This crossing has not been observed in other SARS-CoV-2 spike structures except in XBB.1.5 (**Extended Data Fig. 8c**), where N2 was stabilized by an NTD-directed monoclonal antibody^49,50^. In contrast, BA.3.2.1 showed poorly resolved N1 and N2 (**Extended Data Fig. 8d**). Structural modeling suggested that BA.3.2.1 adopted a shorter N2 with N1 positioned outside N2, akin to WA1 (PDB: 6WR8) (**Fig. 3b, Extended Data Fig. 8e**).

**Fig. 3:**
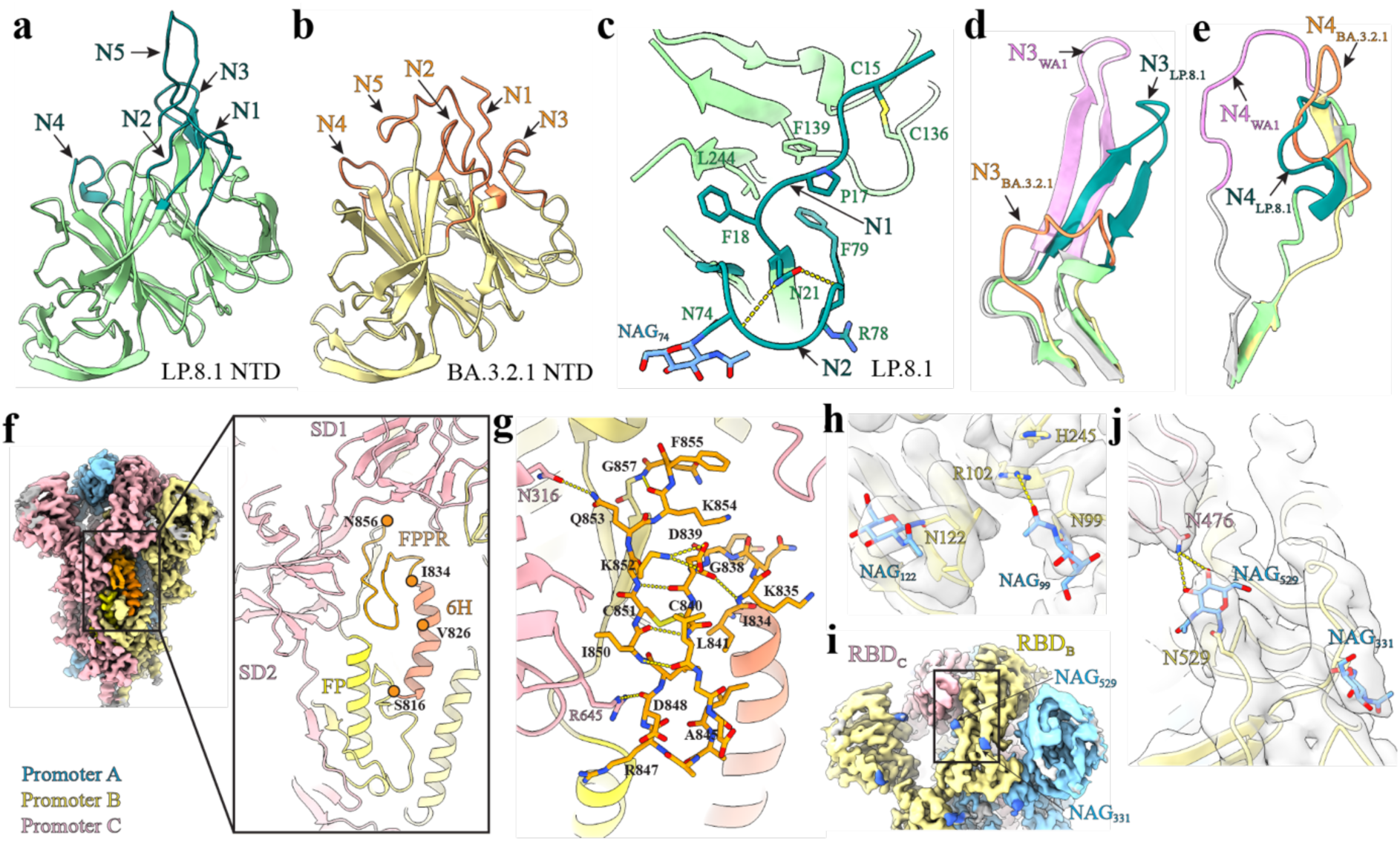
Structural analysis of BA.3.2.1 spike. **a-b**, Alphfold/Cryo-EM model of LP.8.1 NTD (**a**) and BA.3.2.1 NTD (**b**). LP.8.1 loops are colored teal and BA.3.2.1 loops orange. **c,** Interaction network stabilizing the thread-through arrangement of N1 and N2 loops in LP.8.1 NTD. **d-e,** Magnified view of the N3 (**d**) and N4 loops (**e**) in WA1 (plum), BA.3.2.1 (orange), and LP.8.1 (teal) NTDs. **f,** Structural model of BA.3.2.1 FPPR. Left, side view of the cryo-EM map of the closed BA.3.2.1 spike with the FPPR boxed in black; Right, magnified view of the boxed region. FP, FPPR, and 6H are colored yellow, orange, and salmon, respectively. **g,** Interaction network stabilizing the BA.3.2.1 FPPR. Dashed lines represent hydrogen bonds or salt bridges. **h**, Magnified view around the N99 glycan site in BA.3.2.1 spike with the cryo-EM density in gray. **i**, Cryo-EM map around the N529 glycan in protomer B of the BA.3.2.1 spike. **j,** Magnified view of the N529 glycan site in BA.3.2.1 with the cryo-EM density in gray.

BA.3.2.1 carried a rare Δ136-147 that markedly remodels N3, replacing its original β-sheets with a shortened loop (**Fig. 3d, Extended Data Fig. 8a**) and a Δ242-243 deletion that shortened and rearranged N5, shifting away from N3 (**Extended Data Fig. 8f**). Correspondingly, N1, N2, and N5 reorganized and changed the antigenic surface, reshaping the NTD-1 supersite. The N3 with Δ136-147, previously reported in BA.2.87.1 but structurally unresolved^51,52^, was characterized here at high resolution (**Fig. 3d**), providing direct structural insight into NTD-1 remodeling. Additionally, BA.3.2.1 S172F and K187T substitutions reshaped N4, yielding an orientation distinct from WA1 (**Fig. 3e**). Together, these features demonstrated a substantial structural variation in the BA.3.2.1 NTD.

### BA.3.2.1 spike adopts a distinct FPPR conformation

In addition, we identified a fusion peptide proximal region (FPPR; residues 834-856) with a distinct conformation in BA.3.2.1 protomer B, despite the absence of FPPR density in all LP.8.1 maps and in most BA.3.2.1 reconstructions (**Fig. 3f, Extended Data Fig 9a-c**). In BA.3.2.1 protomer C, a partially resolved FPPR formed a helical segment resembling previously reported loop-helix-loop configurations (6XM0 chain B; 7TNW chain B^53^) that approached toward SD1 (**Extended Data Fig. 9a, d-e**). In contrast, the well-resolved FPPR in BA.3.2.1 protomer B adopted a loop-only architecture displaced from SD1 (**Extended Data Fig. 3f**). Residues D839-A845 ran approximately parallel to R847-F855, stabilized by a Cys840-Cys851 disulfide bond, a D839-K852 salt bridge, and backbone hydrogen-bond network (**Fig. 3g**). Additional contacts with neighboring regions, e.g., K835/G838, D848/R645, and Q853/N316, further reinforced this configuration. Notably, residues 826-833, which formed a flexible loop in 6XM0 and 7TNW (**Extended Data Fig. 9d-e**), adopted α-helical turns in BA.3.2.1, extending the three-helix (3H) segment^54^ into a six-helix (here termed 6H) spanning 816-833 (**Fig. 3f-g**). Together, these findings highlight pronounced plasticity of the FPPR and its upstream elements, suggesting that local remodeling of the S2 fusion loop may accommodate the tightly closed conformation of BA.3.2.1 spikes and modulate S2 refolding during membrane fusion.

### N-glycosylation features of BA.3.2.1 Spike

The SARS-CoV-2 spike is heavily glycosylated, carrying 22 conserved N-linked glycosylation sites per spiker protomer in the ancestral strain^55^. Spike glycans played critical roles in protein folding, conformational dynamics, receptor binding, and immune evasion^56,57^. BA.3.2.1 and LP.8.1 spikes shared most core N-glycosylation sites but differed in additional patterns (**Extended Data Fig. 10a**). BA.3.2.1 spike acquired three new sites, including N99 (introduced by I101T), N185 (K187T), and N529 (K529N). Our cryo-EM maps detected glycans at N99 and N529. The N99 glycan formed an H-bond with R102, while Δ242-243 enabled H245 to π-stack with R102, reinforcing local stability (**Fig. 3h**). The N529 glycan projected to contact residue N477 on an adjacent down-RBD, strengthening inter-protomer contacts and likely reducing RBD mobility (**Fig. 3i-j**).

In contrast, LP.8.1 lost N17 (due to T19I) but gained N31 (ΔS31), N188 (R190S), and N354 (K356T) (**Extended Data Fig. 10a**). Glycans at N31 and N188 were not resolved in our dataset, reflecting local flexibility, whereas N354 was clearly visualized (**Extended Data Fig. 10b**). N354 glycosylation, conserved in BA.2.86 lineages, has been shown to stabilize the closed state and mask nearby epitopes, thereby reducing ACE2 accessibility and enhancing immune evasion^57^. Notably, the BA.3.2.2 sublineage independently acquired the K356T, enabling N354 glycosylation and indicating convergent evolution at this position.

### Less ACE2 binding to BA.3.2.1 spike than LP.8.1 spike

To define the functional consequences of the structural differences between BA.3.2.1 and LP.8.1 spikes, we compared their binding affinities to hACE2. Biolayer interferometry (BLI) revealed that BA.3.2.1 spike exhibited lower association (*k*_on_; 2.2-fold) and dissociation (*k*_off_; 1.6-fold) rate constants compared to LP.8.1 spike, resulting in a 1.4-fold reduction in hACE2 binding affinity (*K*_D_) (**Fig. 4a & Extended Data Fig. 11a-b**). To assess the impact of spike variants on viral attachment, we measured the attachment efficiency of BA.3.2.1-spike and LP.8.1-spike mNG SARS-CoV-2 on Vero E6 cells at 4 °C. Despite equivalent input, BA.3.2.1-spike virions showed significantly lower attachment than LP.8.1-spike virions (**Fig. 4b**).

**Fig. 4:**
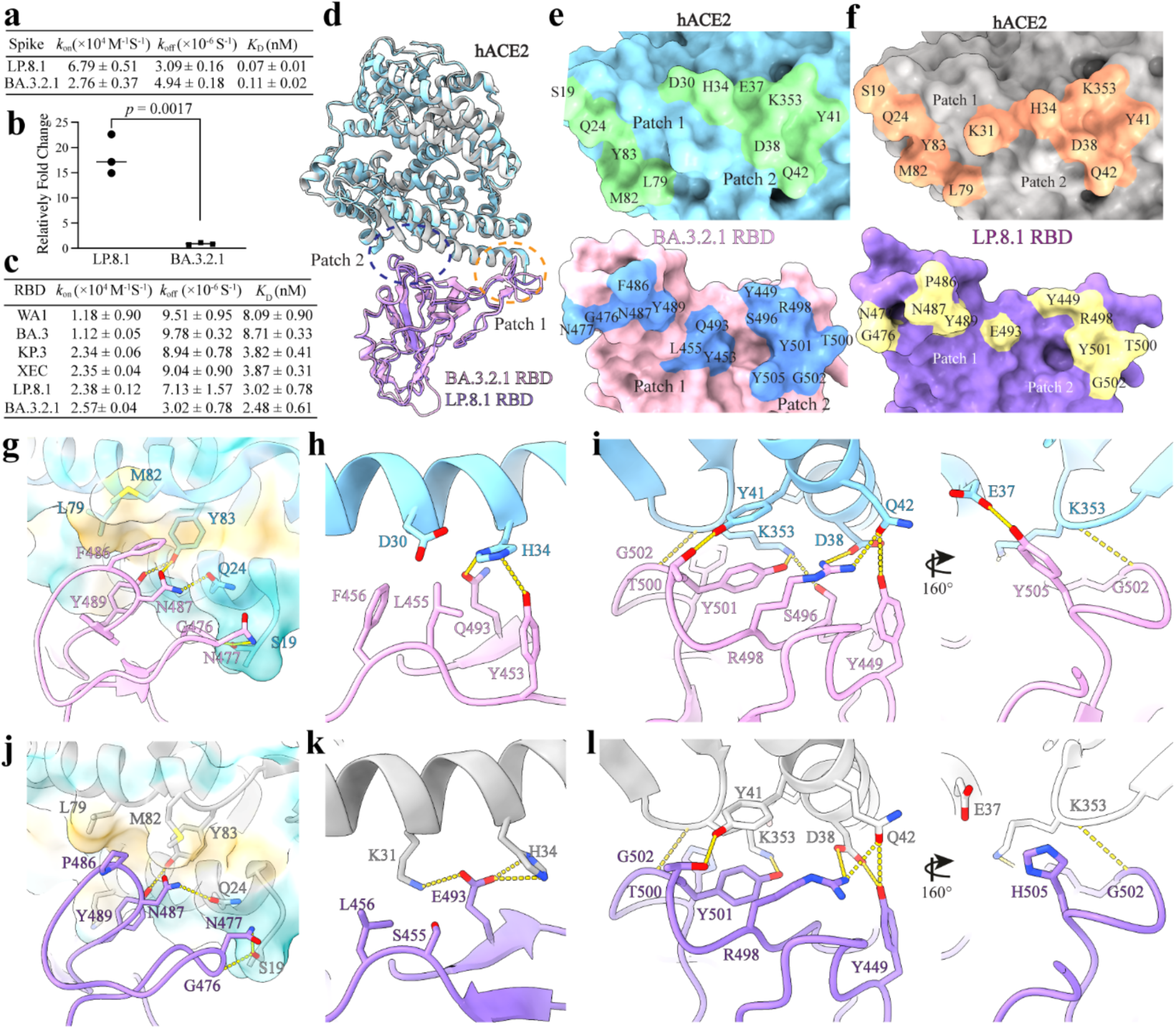
Interaction of BA.3.2.1 and LP.8.1 spike with hACE2. **a,** Summary of BLI kinetics (*k*_on_, *k*_off_, and *K*_D_) for trimeric LP.8.1 and BA.3.2.1 spike binding hACE2. Means± standard deviations variations from three independent experiments are shown. **b,** Quantification of Vero E6 cell-bound BA.3.2.1 and LP.8.1 virus by RT-qPCR. *P* value from a two-tailed unpaired Student’s t-test is shown. **c,** Summary of BLI kinetics (*k*_on_, *k*_off_, and *K*_D_) between LP.8.1 and BA.3.2.1 RBDs binding hACE2. Means± standard deviations variations from three independent experiments are shown. **d,** Structural alignment of hACE2/BA.3.2.1 RBD and hACE2/LP.8.1 RBD complexes. Patches 1 and 2 are indicated. **e-f**, Residue-level views of the RBM-hACE2 interface for BA.3.2.1 (**e**) and LP.8.1 (**f**) (Top, hACE2; bottom, RBD). **g-h**, Interaction networks mediating BA.3.2.1 RBD-hACE2 binding. **j-l,** Interaction networks mediating LP.8.1 RBD-hACE2 binding. In **g** and **j**, the hydrophobic hACE2 surface (L79, M82, Y83) is highlighted. Complexes are shown as cartoons; contact residues are shown as sticks. Dashed lines denote hydrogen bonds or salt bridges; hydrophobic contacts are indicated by residue proximity.

To determine whether ACE2-spike binding differences were attributable to the RBD, we measured hACE2-binding affinities by BLI for recombinant RBDs from BA.3.2.1, LP.8.1, and earlier Omicron sublineages (BA.3, KP.3, XEC), along with the parental WA1 strain. The binding affinity (*K*_D_) ranked as follows: BA.3.2.1 > LP.8.1 > XEC ≈ KP.3 > WA1 > BA.3 **(Fig. 4c & Extended Data Fig. 11c-h)**. Kinetic analysis showed that BA.3.2.1 RBD exhibited a *k*_on_ comparable to JN.1 descendants (KP.3, XEC, and LP.8.1) but a markedly slower *k*_off_, accounting for its stronger affinity. Notably, this high RBD-hACE2 affinity contrasted with the reduced hACE2 binding observed for trimeric BA.3.2.1 spike, consistent with restricted RBD accessibility in a more compact, closed trimer.

To test this accessibility model, we removed the N529 glycan by introducing N529Q into the BA.3.2.1 spike, disrupting the glycan-mediated inter-protomer contact (**Fig. 3j**). BLI showed that BA.3.2.1 N529Q spike displayed a 1.5-fold faster *k*ₒₙ with no detectable change in *k*_off_, yielding a lower *K*_D_ and overall hACE2 binding comparable to LP.8.1 spike (**Extended Data Fig. 11b, i**). These results supported the interpretation that reduced ACE2 binding of BA.3.2.1 arises from limited RBD accessibility imposed by its closed trimeric conformation.

### Cryo-EM structures of BA.3.2.1 and LP.8.1 spike complexed with hACE2

BA.3.2.1 differed from LP.8.1 by >75 spike mutations, many within the RBD (**Extended Data Fig. 1**). To assess whether these changes alter the hACE2-binding interface, we determined cryo-EM structures of each spike in complex with hACE2.

In our initial cryo-EM experiment, incubation of BA.3.2.1 spike with hACE2 at a 1:2 molar ratio on ice did not yield hACE2-bound complexes. The observed particles adopted apo-like conformations, including both closed (3-RBD-down) and 1-RBD-flexible states (**Extended Data Fig. 12a-c**). To increase RBD accessibility, BA.3.2.1 spike was incubated with hACE2 at a 1:3 molar ratio at 37°C, followed by a brief chill on ice prior to plunge-freezing. Under these conditions, nearly all spike particles were hACE2-bound (**Extended Data Fig. 12d-e**). Subsequent high resolution cryo-EM analysis revealed four BA.3.2.1 spike-hACE2 conformations (**Extended Data Fig. 13a-g, Supplementary Table 5**): conformations 1-3 (account for 42.0%, 37.7%, and 13.5% of total particles, respectively) featured 3-RBD-up with one to three hACE2 bound; conformation 4 (account for 6.8% of total particles), displayed a 2-RBD-up arrangement with a single hACE2 bound to an up-RBD (**Extended Data Fig. 13d-g**). By contrast, LP.8.1spike-hACE2 complexes were obtained on ice without elevated temperature (**Extended Data Fig. 14a-c, Supplementary Table 6**). Approximately half of LP.8.1 spikes adopted a 2-RBD-up configuration, with one to two hACE2 bound to up-RBDs, while the remainder were unbound and resembled the apo LP.8.1 conformations (**Extended Data Fig. 14d-e**).

To define the binding interface, we performed local refinement of the BA.3.2.1 RBD-hACE2 and LP.8.1 RBD-hACE2 regions, yielding maps with 3.6 Å and 3.7 Å resolution, respectively (**Extended Data Fig. 13h-i, 14f-g, Supplementary Table 9-10**). Structural alignment indicated overall high similarity between both interfaces, with Cα RMSD of 0.868 Å (**Fig. 4d**). As in the parental and earlier variants, both RBDs engaged hACE2 via two contact patches spanning residues 437-451 (Patch 1) and 498-507 (Patch 2). However, Receptor binding motif (RBM) footprints differed between BA.3.2.1 and LP.8.1 (**Fig. 4e-f**).

Within the BA.3.2.1 RBM-hACE2 interface, patch 1 residues (G476, N477, F486, N487, Y489, L455, Q493, Y453) contacted hACE2 residues S19, Q24, D30, H34, L79, M82, and Y83, whereas patch 2 residues (Y449, S496, R498, T500, Y501, G502, Y505) engaged hACE2 residues E37, D38, Y41, Q42, and K353 (**Fig. 4e**). Relative to WA1 strain, BA.3.2.1 retained multiple conserved contacts, including Y449-Q42 (hACE2), Y453-H34 (hACE2), L455-D30 (hACE2), F486-L79/M82/Y83 (hACE2), Y489-Y83 (hACE2), Q493-H34 (hACE2), T500-Y41 (hACE2), and G502-K353 (hACE2) (**Fig. 4g-i**). As a BA.3 descendant, BA.3.2.1 preserved the cornerstone epistatic pair R498 (interacting with hACE2 D38 and Q42) and Y501 (interacting with hACE2 K353), as well as N477 (interacting with hACE2 S19), all of which strengthened hACE2 binding. Notably, BA.3.2.1 did not carry E493, previously reported to impair hACE2 binding^58^; and gained additional contact with hACE2 K353 via S496 (**Fig. 4i**). Distinct from BA.3, BA.3.2.1 retained Y505-E37 (hACE2) interaction, as in WA1 (**Fig. 4i**).

Compared with BA.3.2.1, LP.8.1 RBD engaged hACE2 through fewer residues in patch 1 (G476, N477, N487, Y489, E493) and in patch 2 (Y449, R498, T500, Y501, G502, H505). Multiple hACE2-binding essential residues (Y449, G476, N477, N487, Y489, R498, T500, Y501, G502) were shared between BA.3.2.1 and LP.8.1 (**Fig. 4f**). Key differences included P486 in LP.8.1 forming hydrophobic contacts with hACE2 L79, M82, and Y83 (**Fig. 4j**), and the signature epistatic “Slip” mutations (L455S+F456L) that allowed E493 to project toward hACE2 and form a strong salt bridge with hACE2 K31 (and an interaction with H34), thereby compensating for the otherwise deleterious effect of E493 on hACE2 binding (**Fig. 4k**). In addition, LP.8.1 lost two contacts with hACE2 K353 and E37 due to G496 and H505, respectively (**Fig. 4l**). Collectively, these results indicated that BA.3.2.1 and LP.8.1 share an overall conserved RBD-hACE2 architecture but implement distinct RBM footprints and interaction networks.

### Structural basis for BA.3.2.1 spike’s antigenicity shift

In line with BA.3.2’s extensive divergence, our neutralization assays (**Fig. 1e-f**) and multiple independent previous reports indicated a marked antigenicity shift of this lineage^12,18,19,38^. To define the structural underpinnings, we analyzed BA.3.2.1 spike models and mapped substitutions across spike subdomains targeted by NAbs (**Fig. 5a**). As in prior variants, most potent NAbs are RBD-directed^26^. BA.3.2.1 encodes 23 RBD substitutions spanning the footprints of five classes of canonical RBD-directing NAbs, thereby recontouring paratope-epitope complementarity (**Fig. 5b-f**).

**Fig. 5:**
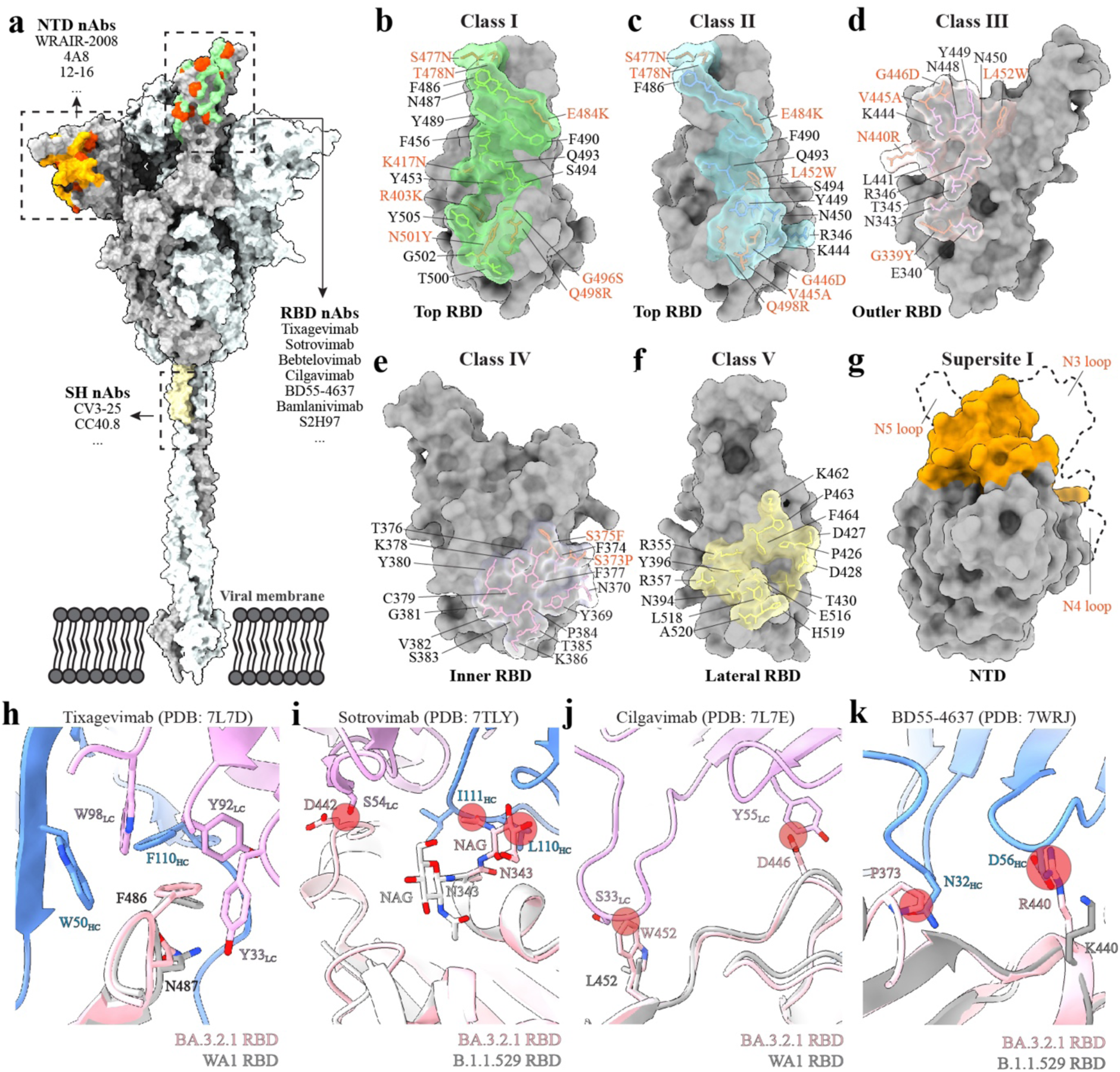
Structural basis for altered antigenic characteristics of BA.3.2.1 spike. **a**, Representative NAbs targeting NTD, RBD, and S2 stem-helix (SH) regions mapped onto a full-length BA.3.2.1 spike in open conformation. Footprints of NTD supersite-1, RBD Classes I-V, and SH targeting NAbs are colored orange, green, and wheat, respectively. BA.3.2.1 mutations are highlighted in red. The full-length BA.3.2.1 spike model was built as previously reported^87^. **b-f**, Primary epitopes for Class I (**b**), Class II (**c**), Class III (**d**), Class IV (**e**), and Class V (**f**) RBD-directed NAbs. The surface representation of BA 3.2.1 RBD is shown in gray. Antibody footprints are rendered as colored surfaces; epitope residues are shown as sticks. BA.3.2.1 substitutions within each epitope are highlighted in red-orange. **g**, Surface representation of the BA.3.2.1 NTD, with supersite-1 highlighted in orange. The dashed outline indicates the corresponding supersite-1 region on WA1 NTD, illustrating extensive reshaping of this neutralizing epitope in BA.3.2.1. **h-k**, Zoomed-in views of the BA.3.2.1 RBD superimposed on reference complexes: WA1 RBD-Tixagevimab (**h**), B.1.1.529 RBD-Sotrovimab (**i**), WA1 RBD-Cilgavimab (**j**), or B.1.1.529 RBD-BD55-4637 (**k**). BA.3.2.1 residues predicted to interfere with antibody binding, and the corresponding antibody contact residues, are shown as sticks. Steric clashes are circled in red. Antibody heavy and light chains are colored blue and plum, respectively.

Class I binds to ACE2-ridge of up-RBD with footprints containing K417, L455-F456, N487, Y489, Q493, Q498, N501, Y505^33^. BA.3.2.1 encodes R403K, K417N, S477N, T478N, E484K, G496S, Q498R, and N501Y within or adjacent to this footprint (**Fig. 5b**). The overall impact of these substitutions can vary by the exact epitope and angle of approach^12^. Several are known to reduce Class I NAb potency. In particular, the K417-E484-N501 triad represented a shared immune escape strategy among SARS-CoV-2 variants^59^. Notably, Tixagevimab (AZD8895) remained active against BA.3.2.1^18^. Structural superposition of the WA1 RBD/Tixagevimab onto the hACE2-bound BA.3.2.1 RBD showed that the key RBM F486/N487 contacts are preserved, maintaining the “aromatic cage” seen with WA1. Specifically, F486/N487 packed against Tixagevimab H-chain W50/F110, and L-chain Y33/Y92/W98, akin to WA1 (**Fig. 5h**). Moreover, BA.3.2.1 lacks Q493E/R, avoiding electrostatic repulsion at the interface (**Extended Data Fig. 15a**). These features rationalized the greater Tixagvimab-sensitivity of BA.3.2.1 than BA.3 (carrying Q493R)^18^.

Class II binds to lateral RBM with footprints including E484, F486, Q493, S494, G496, as well as L452 and S477/T478^26,60^. BA.3.2.1 substitutions G446D, L452W, S477N, T478N, E484K, and Q498R clustered at this interface (**Fig. 5c**), and are expected to impair Class II NAb binding via electrostatic rewiring (G446D, E484K, and Q498R) and steric bulk (L452W). For example, E484K could ablate bamlanivimab (LY-CoV555) by disrupting central RBM contacts (**Extended Data Fig. 15b**).

Class III binds to the outer RBD face, often at or near the N343-glycan patch and the N440-K444-V445-G446 loop region^26,31^. This class is sensitive to loop geometry and electrostatic shifts (e.g., N440K/R and G446D/S/V)^61^. BA.3.2.1 carries G339Y, N440R, V445A, G446D, and remodels the 438-451 and 497-505 loops (**Fig. 5d, Extended Data Fig. 7a**). These changes could (i) penalize Sotrovimab by perturbing glycan-environment and introducing steric clashes (**Fig. 5i, Extended Data Fig. 15c**), (ii) reduce Bebtelovimab complementarity via V445A/G446D and loop shifts (**Extended Data Fig. 15c-d**), and (iii) disrupt Cilgavimab binding through clashes (W452-S33/N34 and D446-Y55) in the RBM-shoulder (**Fig. 5j**). These structural insights accord with abolished activity for Sotrovimab, Bebtelovimab, and Cilgavimab reported against BA.3.2^18^.

Class IV (e.g. BD55-4637) targets a largely conserved cryptic site near the RBD base (e.g., K378/Y380/V382/L390/F392/T430), accessible in up-RBDs^26,62^. BA.3.2.1 S373P within the 371-375 loop remodels the epitope and confers resistance to BD55-4637 by introducing steric clashes between P373-N32, as well as between K440-D56 (**Fig. 5e, k**), explaining the loss of neutralization for BD-55-4637^12,18^. Class V (e.g., S2H97) targets a highly conserved, cryptic region on the lateral RBD^33,63^ (**Fig. 5f**). BA.3.2.1 retains key residues in this region (**Extended Data Fig. 15e**), suggesting preserved sensitivity to S2H97. Notably, the closed BA.3.2.1 spike may occlude these cryptic sites and reduce neutralization of Class IV and Class V NAbs.

Prior work identified NTD-directed NAbs, including 4A8 and WRAIR-2008 targeting antigenic supersite (N1-N5 loops) and 12-16 targeting NTD-SD1 interface^37,64,65^. BA.3.2.1 NTD contains Δ136-147, Δ211, Δ243-244, and additional substitutions that substantially reshape the NTD, such as N3 and N4 loops, and likely ablate these antibodies (**Fig. 3b, 5g, Extended Data Fig. 15f-h**). Despite the absence of EM density for S2 in our maps, the conserved linear stem-helix epitope (residues 1140-1167) targeted by CV3-25 and CC40.8^35,66^ showed no sequence change in BA.3.2.1 (**Extended Data Fig. 1**), and these antibodies were therefore expected to be largely unaffected.

Collectively, our analyses indicate that BA.3.2.1 recontours RBD/NTD epitopes. In concert with a closed trimer that occludes cryptic/shoulder sites, these changes mediate broad immune escape and account for the observed antigenicity shift.

## Discussion

BA.3.2 represents a further evolutionary leap of SARS-CoV-2 with antigenic and functional properties distinct from contemporaneous JN.1 descendants. Using the BA.3.2.1 sublineage as an experimental model, this study delineates how its heavily mutated spike modulates replication, hACE2 engagement, and immune escape. Cryo-EM structures of the apo BA.3.2.1 spike and its ACE2-bound complexes reveal distinctive structural features that mechanistically rationalize immune escape coupled to attenuated replication fitness, a trade-off that likely has constrained BA.3.2 prevalence relative to co-circulating JN.1 descendants.

BA.3.2.1 spike differs from its parental BA.3 by 39 amino acids and from JN.1 descendants by >80 amino acids. Functionally, BA.3.2.1 spike replicates more slowly than JN.1 derivatives in Calu-3 cells and human airway epithelial (HAE) cultures (**Fig. 1**). Compared with BA.3, BA.3.2.1 shows slightly reduced replication in Calu-3 cells, but yet replicates more robustly in HAE, underscoring the value and robustness of paired-competition assays in physiologically relevant systems. Nevertheless, our data align with the low global prevalence of BA.3.2 sublineages detected since late 2024. Notably, replication capacity among JN.1 descendants (XEC>LP.8.1; XEC>KP.3) does not mirror their epidemiological succession (KP.3→XEC→LP.8.1), indicating that variant success reflects factors beyond spike-encoded replication properties alone, e.g., non-spike mutations, host population immunity, and ecological context.

Our results clarify the contribution of S1/S2 cleavage to fitness. Proteolytic priming at residues 685-686 is a critical activation step, and perturbations at these sites can affect syncytia formation, lung pathogenesis, and transmission efficiency^67–69^, however, cleavage efficiency did not predict replication or epidemiologic success across the variants tested: KP.3, BA.3.2.1, and WA1 spikes were efficiently cleaved, whereas BA.3, XEC, and LP.8.1 were only partially cleaved (**Fig. 1b-c**). Together with prior work^40^, these observations argue that enhanced S1/S2 cleavage alone is insufficient to improve fitness. Instead, replication capacity reflects a composite of multiple factors, e.g., spike biophysical properties, receptor engagement, entry pathway usage, innate-immune antagonism, and non-spike genetic background, while population-level immune selection further shapes variant prevalence.

A central insight from our structural analyses is that BA.3.2.1 assembles a compact, “locked” spike trimer, which coherently explains its phenotypes. Relative to LP.8.1, BA.3.2.1 trimers display reduced conformational heterogeneity with the majority adopting the all-RBD-down state (**Fig. 2a-b**). Vector-based quaternary analysis demonstrates compact intra-protomer hinging, a distinctive SD1/SD2 auxiliary configuration, and contracted inter-RBD spacing, together with additional inter-RBD contacts and loop remodeling in 438-450 and 497-506 that likely stabilize the closed conformation (**Fig. 2e-k**). Importantly, K529N introduces an N-glycan at N529 that contacts N477 on the adjacent RBD, reinforcing trimer closure (**Fig. 3h-i**).

This closed, compact architecture has three major functional consequences. First, it reduces trimer breathing and increases thermal stability, consistent with the higher melting temperature of BA.3.2.1 relative to LP.8.1 and its lower structural heterogeneity (**Fig. 2a-c**). Enhanced thermostability may aid persistence of viral infectivity in nature and thereby indirectly influence transmission. Second, the closed state limits RBD up-down transitions, thereby reducing ACE2 accessibility at the trimer level. Consistently, trimeric BA.3.2.1 binds hACE2 less than LP.8.1 despite higher monomeric RBD affinity (**Fig. 3a-c**), and removing the N529 glycan (N529Q) improves trimer-ACE2 binding (**Extended Data Fig. 11i**). Temperature-dependent ACE2-spike complex formation (**Extended Data Fig. 12**) further supports a model in which conformational gating limits receptor engagement, contributing to observed attenuated replication phenotypes relative to JN.1 descendants (**Fig. 1d-h**). The disconnect between intrinsic RBD affinity and whole-spike receptor access has been observed across Omicron lineages and helps rationalize paradoxical “high-escape/low-entry” phenotypes^70^. Third, a closed trimer augments immune evasion, a recurrent strategy among SARS-CoV-2 variants^12,71–73^. The all-RBD-down ensemble sterically occludes neutralizing surfaces, especially cryptic/shoulder epitopes whose recognition depends on RBD opening (**Fig. 5a-f**). Together with extensive RBD and NTD remodeling, this dual mechanism explains the marked antigenicity shift and broad neutralization escape of BA.3.2 observed here (**Fig. 1i-j**) and by others^12,18,38^.

Despite reduced trimer-level ACE2 engagement, BA.3.2.1 preserves most conserved ACE2 contacts at the RBM (**Fig. 4d-i**). We posit that this reconfigured interaction network, including the R498-Y501 epistatic pair with hACE2 D38/Q42 and K353, S496-K353 (hACE2) contact, and Y505-E37 (hACE2), allowing high intrinsic RBD affinity while maintaining strong immune escaping capacity resulted from extensive mutations. The BA.3.2.1 spike architecture effectively decouples intrinsic RBD affinity from trimer accessibility, thus providing a mechanistic basis for the combination of strong antibody escape and attenuated replication. In the pre-attachment state, the spike trimer favors a tightly closed conformation that limits premature exposure of antibody-targeted epitopes; upon receptor engagement, the BA.3.2.1 RBD binds hACE2 tightly to secure efficient viral entry.

Beyond the RBD, BA.3.2.1 exhibits substantial NTD remodeling. The NTD contains a major antigenic supersite targeted by many NTD-directed neutralizing antibodies^74,75^ and also supports many other critical functions, such as NTD-RBD crosstalk, sialic-acid binding, and entry/fusogenicity modulation^76–78^. BA.3.2.1 harbors Δ136-147 (N3) and Δ243-244 (N5) together with additional substitutions that recontour the supersite and distinguish BA.3.2.1 from LP.8.1(**Fig. 3a-e**), likely contributing to escape from the supersite-targeted NAbs in concert with RBD-mediated immune evasion (**Extended Data Fig. S15**). In addition, NTD remodeling may accommodate the hyper-closed quaternary architecture (**Fig. 2f-g**). These observations also raise the hypothesis that NTD remodeling may partially compensate for limited RBD accessibility during early attachment (e.g., glycan-mediated interactions), warranting further investigation.

The limited global expansion of BA.3.2 to date is consistent with WHO risk assessments and independent epidemiological updates: BA.3.2 has remained a Variant Under Monitoring with no sustained growth advantage over co-circulating lineages, despite pronounced antigenic drift^17^. Our data provides a structural-functional basis for this observation. On the vaccine front, human studies indicate that LP.8.1-matched monovalent formulations elicit robust neutralization of JN.1 descendants (including more recent XFG) with expected reduction relative to BA.3.2^79^, aligning with global guidance that JN.1 derivatives’ antigens remain appropriate. However, BA.3.2-like antigenic distinct strains can potentially challenge breadth if they acquire compensatory mutations.

Our study has limitations. The use of an mNG SARS-CoV-2 backbone does not capture non-spike contributions to fitness nor potential spike/non-spike epistasis. Nevertheless, our phenotype-structure concordance and external reports showing comparable replication of BA.3.2 isolates with LP.8.1 in cell culture suggest that spike is a major determinant of the observed differences, while not exclusive^38^. Our serum cohorts were derived from KP.2-/KP.3-infections in a single geography and timeframe. While other reports using panels with diverse immune backgrounds support our observations^12,18^, factors such as time since infection, prior vaccination/boosting, and hybrid-immunity heterogeneity can influence neutralization breadth. We did not evaluate mucosal immunity or T-cell responses, both of which shape protection independent of neutralization titers. Site-specific glycoproteomics (including occupancy and microheterogeneity) will be valuable to study the glycan function in epitope masking or conformational equilibria. Moreover, while our cryo-EM captures multiple apo and ACE2-bound states, time-resolved approaches will be helpful to quantify opening kinetics and membrane-fusion efficiency. Finally, we identify remodeling of the fusion peptide proximal region (FPPR) and an extended 3-helix (3H) segment in BA.3.2.1, but their functional significance during membrane fusion remains to be defined.

In summary, this study reveals that BA.3.2.1 achieves broad neutralizing-antibody escape through epitope substitutions and structural remodeling across the RBD and NTD, but incurs a fitness cost from a hyper-closed, glycan-reinforced quaternary architecture that restricts RBD accessibility. This trade-off explains the persistent but slow expansion of BA.3.2 relative to co-circulating JN.1 descendants and underscores conformational gating as a key determinant of variant fitness, informing next-generation therapeutics and vaccines. Continued surveillance will be essential to detect future trajectories in which variants rebalance this trade-off through compensatory adaptations.

## Material and Methods

### Ethical statement

The use of human serum specimens in this study was reviewed and approved by the University of Texas Medical Branch (UTMB) Institutional Review Board (IRB number 20-0070). No informed consent was required because these deidentified sera were leftover specimens from the routine standard of care and diagnostics. No diagnosis or treatment was involved either.

All virus infections were conducted in a biosafety level 3 (BSL-3) facility with redundant fans in the biosafety cabinets at UTMB. All personnel wore powered air-purifying respirators (Breathe Easy, 3M) with Tyvek suits, aprons, booties, and double gloves.

### Cells

Vero E6 (ATCC® CRL-1586) cells were obtained from the American Type Culture Collection (ATCC, Bethesda, MD). VeroE6 cells expressing TMPRSS2 (JCRB1819) were purchased from SEKISUI XenoTech, LLC. Both cells were cultured in Dulbecco’s modified Eagle’s medium (DMEM) supplemented with 10% fetal bovine serum (FBS; HyClone Laboratories, South Logan, UT) and 1% penicillin/streptomycin (P/S). All cultures were maintained at 37°C with 5% CO_2_. The 293F Freestyle cells were obtained from ThermoFisher Scientific and maintained in FreeStyle™ 293 Expression Medium at 37°C with 8% CO_2_. Cells were tested Mycoplasma negative by nested PCR and fluorescence microscopy at the Tissue Culture Core Facility (TCCF) at UTMB.

### Human serum

Two panels of human sera collected at UTMB were used in the study. The first sample panel, consisting of 26 sera, was collected 21-99 days (median 42.5) after KP.2 infection (as determined by RT-PCR) from individuals aged 22-82 years (median 38.5) who had received 0-4 doses of mRNA vaccines. The second sample panel, consisting of 21 sera, was collected 23-105 days (median 50) after KP.3 infection (as determined by RT-PCR) from individuals aged 26-96 years old (median 54) who had received 0-6 doses of mRNA vaccines. Patient information was completely deidentified from all specimens. The de-identified human sera were heat-inactivated at 56°C for 30 min before the neutralization test. The serum information is presented in Supplementary Table 1-2.

### Generation of recombinant SARS-CoV-2 mutant viruses

Recombinant BA.3.2.1 (GISAID: EPI_ISL_19771108)-, XEC (EPI_ISL_19283891)-, LP.8.1 (GISAID: EPI_ISL_19674199)-mNG SARS-CoV-2 spike-variants were constructed by engineering the complete *spike* gene from the indicated variants into an infectious cDNA clone of mNG USA-WA1/2020 as reported previously. Briefly, standard overlap PCRs were conducted to introduce the spike mutations into the infectious clone of the mNG USA-WA1/2020. The full-length infectious cDNA clones were assembled by *in vitro* ligation. Subsequently, the genome-length RNAs were synthesized by *in vitro* transcription. The full-length RNA and N gene RNA transcripts were electroporated into Vero E6-TMPRSS2 cells to rescue the recombinant viruses. After 48-72h of transfection, the supernatants (referred to as P0) were harvested. The P0 stocks were then inoculated into freshly prepared Vero E6 cells for further amplification. At 48 hours post-infection, supernatants (P1) were harvested, clarified by centrifugation at 1000 × g for 10 mins, and stored at −80 ℃.

### RNA extraction, RT-PCR, and cDNA sequencing

Cell culture supernatants were mixed with a five-fold excess of TRIzol LS Reagent (ThermoFisher Scientific, Waltham, MA). Viral RNAs were extracted according to the manufacturer’s instructions. The extracted RNAs were dissolved in 50 μL nuclease-free water. For sequence validation of mutant viruses, 10 ng of RNA samples were used for reverse transcription by using the SuperScript IV First-Strand Synthesis System (ThermoFisher Scientific) with CoV-21115V and CoV-YH5 (Supplementary Table 4). The resulting DNAs were purified by the QIAquick Gel Purification Kit and sequenced by Sanger sequencing at GENEWIZ (South Plainfield, NJ).

### Plaque assay

1×10^6^ Vero E6-TMPRSS2 cells per well were seeded into 6-well plates. The next day, 200 μl of 10-fold serially diluted virus was added to pre-seeded cells and incubated at 37°C for 1 hour. After that, the inoculum was replaced with 2 ml of overlay medium containing DMEM with 2% FBS, 1% penicillin/streptomycin, and 1% Seaplaque agarose (Lonza, Walkersville, MD, USA). After 2 days of incubation at 37°C, 2 ml of overlay medium supplemented with neutral red (Sigma-Aldrich, St. Louis, MO, USA) was added to stain the cells, and plaques were counted on the next day.

### Growth kinetics of recombinant SARS-CoV-2 in cell culture

Approximately 3 × 10^5^ Calu-3 cells were seeded into each well of 12-well plates and cultured at 37°C, 5% CO_2_ for 24 h. WA1, BA.3, BA.3.2.1, KP.3, XEC, or LP.8.1 was inoculated into the cells at an MOI of 0.05. The virus was incubated with the cells at 37°C for one hour. After infection, the cells were washed 3 times with DPBS to remove unattached virus. One milliliter of culture medium was added to each well to maintain the cells. At each time point, 200 μL of culture supernatant was collected for the plaque assay. Meanwhile, 200 μL fresh medium was added to each well to replenish the culture volume. The cells were infected with triplicate for each virus. All samples were stored at −80°C until plaque analysis.

### Virion purification and spike protein cleavage analysis

One milliliter of each virus from the P1 stocks collected from Vero E6 cells was combined with polyethylene glycol (PEG)-8000 (Sigma-Aldrich) to achieve a final concentration of 10%. The mixtures were incubated at room temperature for 30 minutes, then centrifuged at 4000 × g for 10 minutes to pellet the virions. The resulting pellets were washed once with 70% ethanol and subsequently resuspended in 2× Laemmli sample buffer (Bio-Rad) containing 0.7 M β-mercaptoethanol (BME). Samples were heat-inactivated at 95°C for 15 minutes, centrifuged at 13,000 × rpm for 10 minutes, and then separated on a 4-15% gradient SDS-PAGE gel. Viral spikes and nucleocapsid proteins were detected by Western blot using specific antibodies: anti-S1 (Sino Biological #40591-T62) and anti-N (Sino Biological #40143-R001), respectively. Densitometric quantification was performed using ImageJ.

### Fluorescent focus reduction neutralization test (FFRNT)

Neutralization titers of human sera were evaluated using FFRNT against BA.3-, BA.3.2.1-, KP.3-, KP.2-, XEC-, and LP.8.1-spike mNG SARS-CoV-2s, following a previously established FFRNT protocol^13^. In brief, 3 × 10^4^ Vero E6 cells were plated per well in 96-well plates (Greiner Bio-one™) and incubated overnight. The following day, sera were serially diluted 2-fold starting at 1:20, covering a final dilution range of 1:20 to 1:20,480. Each diluted serum sample was mixed with 100-150 focus-forming units (FFUs) of mNG SARS-CoV-2 and incubated at 37°C for 1 hour. Afterwards, the mixtures were added to the Vero E6 cell monolayers. After a 1-hour infection period, the inoculum was removed and replaced with 100 μL overlay medium containing 0.8% methylcellulose. Plates were then incubated at 37°C for 16 hours. Fluorescent foci were imaged using a Cytation™ 7 system (BioTek) equipped with a 2.5 × FL Zeiss objective, and images were processed via Gen5 software with settings optimized for GFP detection (wavelengths 469-525 nm, threshold 4000, object size 50-1000 μm). The number of foci per well was quantified and normalized to controls without serum to calculate relative infectivity. The FFRNT_50_ titer was defined as the lowest serum dilution, reducing fluorescent foci by more than 50%. Each serum was tested in duplicate, and the geometric mean was calculated. Data visualization was performed in GraphPad Prism 9, and figures were prepared using Adobe Illustrator. For analysis and plotting, an FFRNT_50_ of <20 was assigned a value of 10.

### Pairwise competition experiment

Human primary airway epithelial (HAE) cultures (EpiAirway) were obtained from MatTek Life Sciences. This 3D mucociliary tissue model is derived from normal human tracheal/bronchial epithelial cells. Two mNG SARS-CoV-2 spike variants were mixed at a 1:1 ratio based on viral titers (PFU/ml), as determined by plaque assays on VeroE6-TMPRSS2 cells. The inoculum was prepared in DPBS, with each virus at a final concentration of 1 × 10⁶ PFU/ml. An aliquot of this virus mixture was stored to confirm the input ratio of the two variants.

Before infection, HAE cultures were incubated with DPBS at 37°C for 30 minutes. Following removal of DPBS, 200 µl of the viral inoculum was applied to the apical surface of each well. After a 2-hour incubation at 37°C with 5% CO₂, the inoculum was removed, and cultures were washed three times with DPBS to eliminate unbound virus. At each collection time point, 300 µl of DPBS was added to the apical surface and incubated at 37°C with 5% CO₂ for 30 minutes to recover released viruses. The DPBS washes were collected into 2-ml tubes. On day 4 post-infection, after collecting the apical washes, 300 µl of TRIzol reagent (Invitrogen) was added to the cells to lyse them and preserve total cellular RNA. All samples were stored at −80°C until further processing.

For analysis, 100 µl of each sample was mixed with Trizol LS reagent (Thermo Fisher Scientific) at a 1:5 ratio. Viral RNA was extracted using the Direct-zol RNA Miniprep Plus kit (Zymo Research) following the manufacturer’s protocol. To identify and differentiate between the variants, cDNA fragments (100-200 bp) containing strain-specific mutations were amplified using the SuperScript™ IV One-Step RT-PCR System with designated primer pairs. Primer set 1: K529N-F and K529N-R targeted *spike* gene residues 499-570, distinguishing BA.3 from BA.3.2.1. Primer set 2: A852K-F and A852K-R amplified residues 823-883 to differentiate BA.3.2.1 from LP.8.1. Primer set 3: K1086R-F and K1086R-R covered residues 1045-1122, distinguishing LP.8.1 from XEC. Primer set 4: F59S-F and F59S-R targeted residues 31-97 to differentiate KP.3 from XEC. Primers are listed in Supplementary Table 4. The resulting cDNA amplicons were purified via gel extraction and submitted for Illumina next-generation sequencing (NGS) at the UTMB sequencing core facility.

### Protein expression and purification

The cDNAs encoding the ectodomain (residues 1-1208) of BA.3.2.1 (GISAID: EPI_ISL_19771108) and LP.8.1 spike (GISAID: EPI_ISL_19674199) with HexaPro (F817P, A892P, A899P, A942P, K986P, and V987P), a mutated fusion cleavage site (_682_GSAS_685_) and a C-terminal T4 fibritin trimerization domain, a strep-tag II, and a 6x His tag, were cloned into the mammalian cell expression vector pαH^24,70^. The BA.3.2.1 spike carrying the N529Q mutation was generated by site-directed mutagenesis using N529Q-F and N529Q-R (Supplementary Table 4) and cloned into the same vector. To express the spike protein, 500 mL FreeStyle 293-F cells were transiently transfected with 0.5 mg of plasmid using Polyethylenimine Hydrochloride (MW 40,000) (Polysciences). Cultures were harvested five days after transfection, and the supernatant was collected by centrifugation. The secreted S protein in the supernatant was passed through a 0.22 μm filter and purified using Ni-NTA agarose columns (Qiagen). Spike protein was further purified by size-exclusion chromatography using a Superose 6 10/300 column (Cytiva) in a buffer composed of 20 mM Tris pH 8.0, 200 mM NaCl.

A gene encoding human ACE2 (1-615) with a C-terminal 6xHis tag was cloned into vector pCAG^80^. The expression vectors were transiently transfected into FreeStyle 293-F cells as described above. Four days after transfection, the supernatant was harvested. The secreted ACE2 protein in the supernatant was passed through a 0.22 μm filter and purified using Ni-NTA agarose columns. ACE2 protein was further purified by size-exclusion chromatography using a Superdex 200 10/300 column (Cytiva) in a buffer composed of 20 mM Tris pH 8.0, 150 mM NaCl.

The RBD (residues R319-F541) of WT, BA.3, BA.3.2.1, KP.3, XEC, and LP.8.1 spike with an N-terminal signal peptide for secretion and a C-terminal 6×His tag for purification was inserted into the vector pCAG^72^. The secreted RBD protein in the supernatant was passed through a 0.22 μm filter and purified using Ni-NTA agarose columns. RBD protein was further purified by size-exclusion chromatography using a Superdex 200 10/300 column in a buffer composed of 20 mM Tris pH 8.0, 150 mM NaCl.

### Cryo-EM sample preparation and imaging

For BA.3.2.1 and LP.8.1 spike alone, a 3 μL sample at 1.3 mg/mL was applied to QUANTIFOIL R1.2/1.3 grid (Electron Microscopy Sciences) that had been plasma-cleaned by PELCO easiGlow™ Glow Discharge Cleaning System (TED PELLA, Inc.). The grid was blotted for 2 seconds and plunge-frozen in liquid ethane using Vitrobot Mark IV (ThermoFisher Scientific) at 8 °C and 100% humidity. For LP.8.1 spike-hACE2 complex, hACE2 was mixed with the spike at a 2:1 molar ratio to a final spike concentration of 1.2 mg/ml and incubated on ice for 30 minutes. 3 µL samples were applied to the grid as described above. For BA.3.2.1 spike-hACE2 complex, hACE2 was mixed with the spike at a molecular ratio of 3 to 1 to a final concentration of spike at 1.2 mg/ml and incubated at 37°C for 15 minutes, followed by incubation on ice for 30 minutes. 3 µL samples were applied to the grid as described above.

Grids were loaded into a Titan Krios G3i microscope (Thermo Fisher Scientific) equipped with a K3 direct electron detector with a GIF Quantum energy filter (20-eV energy slit) (Gatan) and operating at 300 keV. Cryo-EM data were automatically acquired with a K3 camera using SerialEM at a nominal magnification of 105,100× (corresponding to 0.832 Å per pixel) with a nominal defocus range of −0.9 to −2.5 µm. Forty-frame movie stacks were collected over an exposure time of 1.0 s with a total dose of 39.8-39.94 e^−^ Å^−1^. The detailed data collection parameters were summarized in Supplementary Table 5-6.

### Cryo-EM data processing

Collected movie frames were imported into CryoSPARC^81^ for image processing. All reconstructions were performed using C1 symmetry to avoid imposing symmetry-related bias and to capture potential structural heterogeneity. Movie data were motion-corrected using Patch Motion Correction. The contrast transfer function (CTF) was estimated using Patch CTF Estimation. Micrographs with CTF fit worse than 5 Å were excluded.

For BA.3.2.1 apo spike, a total of 4,474,773 particles were selected using Template Picker and extracted with 2.1 × binning (196 pixels) from 12,806 micrographs. 1,411,706 particles were selected after three rounds of two-dimensional (2D) classification. 400,000 particles were used to create four initial three-dimensional (3D) volumes using Ab Initio. Multiple iterative rounds of heterogeneous refinement were performed using all the particles, resulting in 2 conformations: closed, with 3-RBD-down; flexible, with 2-RBD-down and one RBD flexible. Particles were re-extracted without binning (416 pixels) and first refined using homogeneous refinement, followed by cryoSPARC’s implementation of global and local CTF refinement and a final round of non-uniform refinement, yielding 2.6 Å and 3.0 Å reconstruction for closed and flexible conformation, respectively. To further resolve the NTD and RBD of the BA.3.2 apo spike, local refinement was performed in cryoSPARC using the closed conformation. Masks covering RBD_A_/RBD_C_/NTD_B_, RBD_A_/NTD_C_, RBD_B_/NTD_A_, and NTD_B_ were created in UCSF Chimera^82^. Masks were imported into cryoSPARC with dilation and soft padding. Local refinement was performed with particle re-center, yielding 2.9-3.0 Å reconstructions for RBD_A_/RBD_C_/NTD_B_, RBD_A_/NTD_C_, RBD_B_/NTD_A_, and NTD_B_.

For LP.8.1 apo spike, a total of 13,741,340 particles were selected using Template Picker and extracted with 2.1 × binning (196 pixels) from 15,414 micrographs. 3,618,231 particles were selected after three rounds of 2D classification. 400,000 particles were used to create four initial 3D volumes using Ab Initio. Multiple rounds of heterogeneous refinement were performed using all the particles, resulting in 3 conformations: closed, 3-RBD-down; open, 1-RBD-up; flexible, 2-RBD-down, and one RBD flexible. Particles were re-extracted without binning (416 pixels) and first processed using homogeneous refinement, followed by CryoSPARC’s implementation of global and local CTF refinement and a final round of non-uniform refinement, yielding 2.5 Å, 2.5 Å, and 2.6 Å reconstruction for closed, open, and flexible conformation, respectively. To further resolve the NTD and RBD of the LP.8.1 apo spike, local refinement was performed in CryoSPARC using the closed conformation. Masks covering each down RBD and its neighboring NTD were created using UCSF Chimera. Masks were imported into CryoSPARC with dilation and soft padding. Local refinement was performed with re-center, yielding 2.8 Å reconstructions for each RBD and NTD.

For BA.3.2.1 spike/hACE2 complex, a total of 4,539,114 particles were selected using Blob Picker and extracted with 2.2 × binning (220 pixels) from 12,806 micrographs. After three rounds of 2D classification, 40,000 particles were used to create five initial 3D volumes using Ab Initio. All 3D volumes were used as input for three iterative rounds of heterogeneous refinement and 3D classifications using all particles, resulting in 4 conformations: conformation 1, 3-RBD-up, all were bound by hACE2; conformation 2, 3-RBD-up, one of the up-RBD was bound by hACE2; conformation 3, 3-RBD-up, two of the up-RBD was bound by hACE2; conformation 4, 2-RBD-up, one of the up-RBD was bound by hACE2. Particles were re-extracted without binning (440 pixels) and first refined using homogeneous refinement, followed by CryoSPARC’s implementation of global CTF refinement and a final round of non-uniform refinement, yielding 3.3 Å, 3.2 Å, 3.5 Å, and 3.7 Å reconstruction for conformation 1-4, respectively, according to GSFSC at 0.143. To further resolve the binding interface between ACE2 and RBD, local refinement was performed in CryoSPARC using particles from conformational 1. A mask was created using UCSF Chimera that covers the up-RBD and ACE2. The mask was imported into CryoSPARC with dilation and soft padding. Local refinement was performed with particle re-centering, yielding a 3.6 Å reconstruction. The map was further sharpened using DeepEMhancer.

For LP.8.1 spike/hACE2 complex, a total of 3,194,764 particles were selected using Template Picker and extracted with 2.2 × binning (220 pixels) from 8,294 micrographs. 1,140,024 particles were selected after three rounds of 2D classification. 400,000 particles were used to create five initial 3D volumes using Ab Initio. Multiple rounds of heterogeneous refinement were performed on all particles, yielding four conformations. Conformation 1 and 2 are LP.8.1 spike/hACE2 complex: conformation 1, 2-RBD-up, one of the up RBD was bound by hACE2; conformation 2 also has two up RBD, and both are bound by hACE2. Conformation 3 and 4 are apo spike. Conformation 3 has one up RBD, and conformation 4 is in closed conformation. Particles were re-extracted without binning (480 pixels) and first refined using homogeneous refinement, followed by CryoSPARC’s implementation of global and local CTF refinement and a final round of non-uniform refinement, yielding 2.8 Å, 3.8 Å, 2.8 Å, and 3.2 Å reconstruction for conformation 1-4, respectively. To further resolve the RBD-hACE2 interface, local refinement was performed in CryoSPARC using particles from conformation 1. A mask covering the up-RBD with hACE2 binding was created using UCSF Chimera. Masks were imported into CryoSPARC with dilation and soft padding. Local refinement was performed without particle subtraction, yielding a 3.7 Å reconstruction. The map was further sharpened using DeepEMhancer.

Local resolutions of final reconstructions were estimated by CryoSPARC’s blocres and displayed as resolution ranges in UCSF ChimeraX.

### Model building, refinement, and analysis

Cryo-EM structure of the BA.3 spike (PDB: 7XIY) and KP.3.1 spike (PDB: 9ELI, 9ELH) was used to build an initial model of the apo BA.3.2.1 spike and the apo LP.8.1 spike, respectively. The initial model was built by rigid body fitting in UCSF Chimera and manual adjustments in Coot. The model was refined iteratively by real-space refinement in Phenix and by manual adjustments and improvements in Coot. Initial model building for BA.3.2.1 RBD/hACE2 complex, LP.8.1 RBD/hACE2 complex was built by rigid body fitting of BA.3.2.1 RBD, LP.8.1 RBD, and hACE2 in UCSF Chimera and manual adjustments in Coot. The model was refined as described above. The full-length BA.3.2.1 NTD model was built using the Predict and Build tool in Phenix^48^.

For 3D variability analysis, EM maps of BA.3.2.1 and LP.8.1 spikes were separately analyzed in CryoSPARC with resolution filtered at 5 Å. A movie of the component was generated in UCSF Chimera using frames from the 3DVA display program in simple mode in CryoSPARC (Supplementary Movies 1-5)^83^.

Protein-protein interaction was analyzed using UCSF ChimeraX and PISA^84^. Figures and movies were prepared using UCSF ChimeraX. Final map and model statistics are summarized in Supplementary Table 5-6, 9-10.

### Negative-stain Electron Microscopy

BA.3.2.1 spike was incubated with hACE2 at a molar ratio of 1:3 at 37°C for 15 minutes. A 4 µL sample at 0.05 mg/mL was applied to a glow-discharged CF200-Cu carbon film grid (Electron Microscopy Sciences). The sample was left on the carbon film for 60 s, followed by negative staining with 2% uranyl formate for 60 s. Micrographs were recorded on a JEOL 2100FS microscope at 80,000 × magnification, operated at 200 keV. The images were imported and processed in CryoSPARC.

### Vector-based analysis

Vector analysis of intraprotomer and interprotomer domain positions was performed as described^42,43^ using the Visual Molecular Dynamics (VMD)^85^ software package Tcl interface (https://zenodo.org/records/4926233). For each protomer, Cα centroids were determined for the NTD (residue 27-43 and 54-271), NTD’ (residue 44-53 and 272-293), NTD sheet motif (NTDa, residue 116-129 and 169-172), NTD’ residue 276 (NTD’a), RBD (residue 330-443 and 503-528), RBD helix motif (RBDa, residue 403-410), SD1 (residue 323-329 and 529-590), SD1 residue 575 (SD1a), SD2 (residue 294-332, 591-620, 641-691, and 692-696), SD2 residue 671 (SD2a), CD (residue 711-716 and 1072-1121), and S2 sheet motif (S2s, residue 717-727 and 1047-1071). Intraprotomer vectors were calculated between the following within protomer centroids: NTD to NTD’, NTD’ to SD2, SD2 to SD1, SD2 to CD, SD1 to RBD, CD to S2s, NTDa to NTD, and RBD to RBDa. Vector magnitudes, angles, and dihedrals were determined from these vectors and centroids. PCA was performed in R with the vector data centered and scaled. Interprotomer vectors were calculated for the following: NTD’ to NTD’a, NTD’ to SD2, SD2 to SD2a, SD2 to SD1, SD1 to SD1a, and SD1 to NTD’. Angles and dihedrals were determined from these vectors and centroids. Vectors for the RBD to adjacent RBD and RBD to adjacent NTD were calculated using the above RBD, NTD, and RBDa centroids. Vectors were calculated for the following: RBD_A_ to RBD_C_, RBD_C_ to RBD_B_, and RBD_B_ to RBD_A_. Angles and dihedrals were determined from these vectors and centroids. PCA was performed in R with the vector data centered and scaled^86^.

### Circular Dichroism

The thermal stability of BA.3.2 and LP.8.1 spike protein was assessed by circular dichroism JASCO J-815 spectropolarimeter (Jasco). Protein samples were prepared at a final concentration of 0.1 mg/mL in PBS. CD spectra were recorded at 222 nm. A thermal melt was performed from 25°C to 85°C, increasing the temperature in 1°C increments, and the CD signal was monitored at each step after equilibration for 1 s. The resulting melting curve was fitted to determine the protein’s thermal denaturation midpoint (Tm).

### Bio-Layer Interferometry analysis

The recombinant, purified RBD or spike proteins were used to measure affinity with hACE2 on the Octet R8 (Sartorius). For RBD/hACE2 interaction, 2 μg ACE2 was biotinylated with NHS-PEO_4_-Biotin (ThermoFisher) and captured onto the SA sensor (Sartorius) for 300 s. The loaded biosensors were then quenched and dipped into PBS buffer for 60 s to adjust baselines. Subsequently, the biosensors were dipped into serially diluted RBD proteins (100∼3.7 nM) for 300 s to record association kinetics and then dipped into PBS buffer for 600 s to record dissociation kinetics. PBS buffer without RBD was used as a background. For spike/hACE2 interaction, 6 μg of biotinylated hACE2 proteins were captured onto the SA sensor (Sartorius) for 300 s. The loaded biosensors were then quenched and dipped in PBS buffer for 60 s to adjust baselines. Subsequently, the biosensors were dipped into serially diluted spike proteins (50∼1.9 nM) and then dipped into PBS buffer containing 0.5% Tween 20 for 600 s to record dissociation kinetics. All steps were performed at 25°C with shaking. Data were analyzed using the Octet Data Analysis software V13 (Sartorius), which was used to fit the curve using a 1:1 binding model and the global fitting method.

### Virus binding and RT-qPCR quantification

Vero E6 cells were seeded into 96-well plates at 3 × 10^4^ cells per well 24 hours before infection. BA.3.2.1 and LP.8.1 viruses were added at an MOI of 0.1, and cells were incubated at 4 °C for 1 hour to allow viral adsorption. After incubation, cells were washed three times with PBS to remove unbound virus and then lysed directly in TRIzol for total RNA extraction. Viral RNA levels were quantified by RT-qPCR using primers 2019-nCoV_N2-F and 2019-nCoV_N2-R targeting the viral N gene. Cellular actin levels were quantified using primers β-actin-F and β-actin-R as normalization controls. RT-qPCR was performed on the QuantStudio 7 Real-Time PCR instrument (ThermoFisher Scientific). Relative RNA levels were obtained by normalizing the viral RNA to cellular actin RNAs using the 2^ΔCt^ method. Statistical significance was assessed using a two-tailed unpaired t-test. Experiments were performed with three independent biological replicates.

### Statistics

Sample size estimation was based on previously published studies; no statistical methods were used to predefine sample sizes. All collected data were included in the analysis. The experiments were not randomized. Patient information was blinded, and investigators were blinded to sample identities during data collection and/or analysis. Neutralization assays were conducted in duplicate, and all replication attempts were successful. Continuous variables were reported as geometric means with 95% confidence intervals or as medians. For geometric mean titer (GMT) calculations and statistical comparisons, sera with undetectable antibody titers (<20) were assigned a value of 10. Comparison between neutralization titers was performed using the Wilcoxon matched-pairs signed-rank test with GraphPad Prism 10.

For competition experiments, six independent infections were performed per group. Simple linear regression was used to assess the statistical significance of RNA ratios at each indicated time point relative to the input RNA ratio, with absolute *p*-values reported. A *p*-value of <0.05 was considered statistically significant. Figures were assembled using Adobe Illustrator.

For viral growth kinetics experiments, four independent infections were conducted per group. Data were log10-transformed to approximate a normal distribution prior to statistical analysis.

## Data availability

All data supporting the findings have been included in this study. The sequence of SARS-CoV-2 strains can be accessed through GISAID (https://gisaid.org) with the following codes: BA.3 (EPI_ISL_7605591), KP.2 (EPI_ISL_ 19002640), KP.3 (EPI_ISL_19016982), XEC (EPI_ISL_19283891), LP.8.1 (EPI_ISL_19674199) and BA.3.2 (EPI_ISL_19771108). The sequence of SARS-CoV-2 mNG has been deposited in the supplemented information of our previous study. The atomic coordinates and cryo-EM maps of the structure of closed BA.3.2 spike (EMD:73393, PDB:9YSJ), local refinement of RBD_C_/RBD_A_/NTD_B_ in closed BA.3.2 spike (EMD: 73599, PDB: 9YX8), local refinement of RBD_A_/NTD_C_ in closed BA.3.2 spike (EMD: 73597, PDB: 9YX7), local refinement of RBD_B_/NTD_A_ in closed BA.3.2 spike (EMD: 73619, PDB: 9YXX), local refinement of NTD_B_ in closed BA.3.2 spike (EMD: 73795, PDB: 9Z3Y), flexible BA.3.2 spike (EMD:73395, PDB: 9YX6), hACE2/BA.3.2 spike complex (EMD: 73404, EMD: 73405, EMD: 73408, EMD: 73409), local refinement of hACE2/BA.3.2 RBD (EMD: 73426, PDB: 9YSR) closed LP.8.1 spike (EMD: 73394, PDB: 9YSK), local refinement of RBD_A_/NTD_C_ in closed LP.8.1 spike (EMD: 73441, PDB: 9YT4), local refinement of RBD_B_/NTD_A_ in closed LP.8.1 spike (EMD: 73443, PDB: 9YT6), local refinement of RBD_C_/NTD_B_ in closed LP.8.1 spike (EMD: 73444, PDB: 9YT7), open LP.8.1 spike (EMD: 73535, PDB: 9YW0), flexible LP.8.1 spike (EMD: 73396), hACE2/LP.8.1 spike complex (EMD: 73427, EMD: 73428, EMD: 73430, EMD: 73439), local refinement of hACE2/LP.8.1 RBD (EMD: 73429, PDB: 9YSS) generated in this study have been deposited in the Protein Data Bank (www.rcsb.org), and Electron Microscopy Data Bank (www.ebi.ac.uk/emdb/).

## Acknowledgments

We thank Dr. Michael B. Sherman for advice and support with cryo-EM data collection; K.-Y. Wong, J. Perkyns, and G. Lynch for computational support; and the Sealy & Smith Foundation for supporting the Sealy Center for Structural Biology and Molecular Biophysics at the University of Texas Medical Branch (UTMB). We are grateful to Dr. Haiping Hao and the UTMB Molecular Genetics Facility for advice and support with next-generation sequencing (NGS) data collection and analysis. We also thank the participants who contributed serum samples. X.X. was supported by NIH contract HHSN272201400006C, and by awards from the Sealy & Smith Foundation, the Kleberg Foundation, the John S. Dunn Foundation, and the Amon G. Carter Foundation. The funders had no role in study design, data collection, data analysis, the decision to publish, or the preparation of the manuscript.

## Author contributions

Conceptualization, X.X.; Methodology, Y.W., Y.H., Z.C., J.Z., K.Z., P.R., B. L., X.X.; Investigation, Y.W., Y.H., Z.C., J.Z., K.Z., B.L., X.X.; Resources, P.R., P.-Y.S., X.X.; Data Curation, Y.W., Y.H., J.Z., P.R., X.X.; Writing-Original Draft, Y.W., Y.H., X.X.; Writing-Review & Editing, Y.W., Y.H., J.Z., Z.C., P.R., P.-Y. S, B. L., X.X; Supervision, X.X.; Funding Acquisition, P.-Y.S., X.X..

