## Extended data Fig1-15; Supplementary table 1-10 for "Functional and Structural Basis of Omicron BA.3.2.1 Spike"

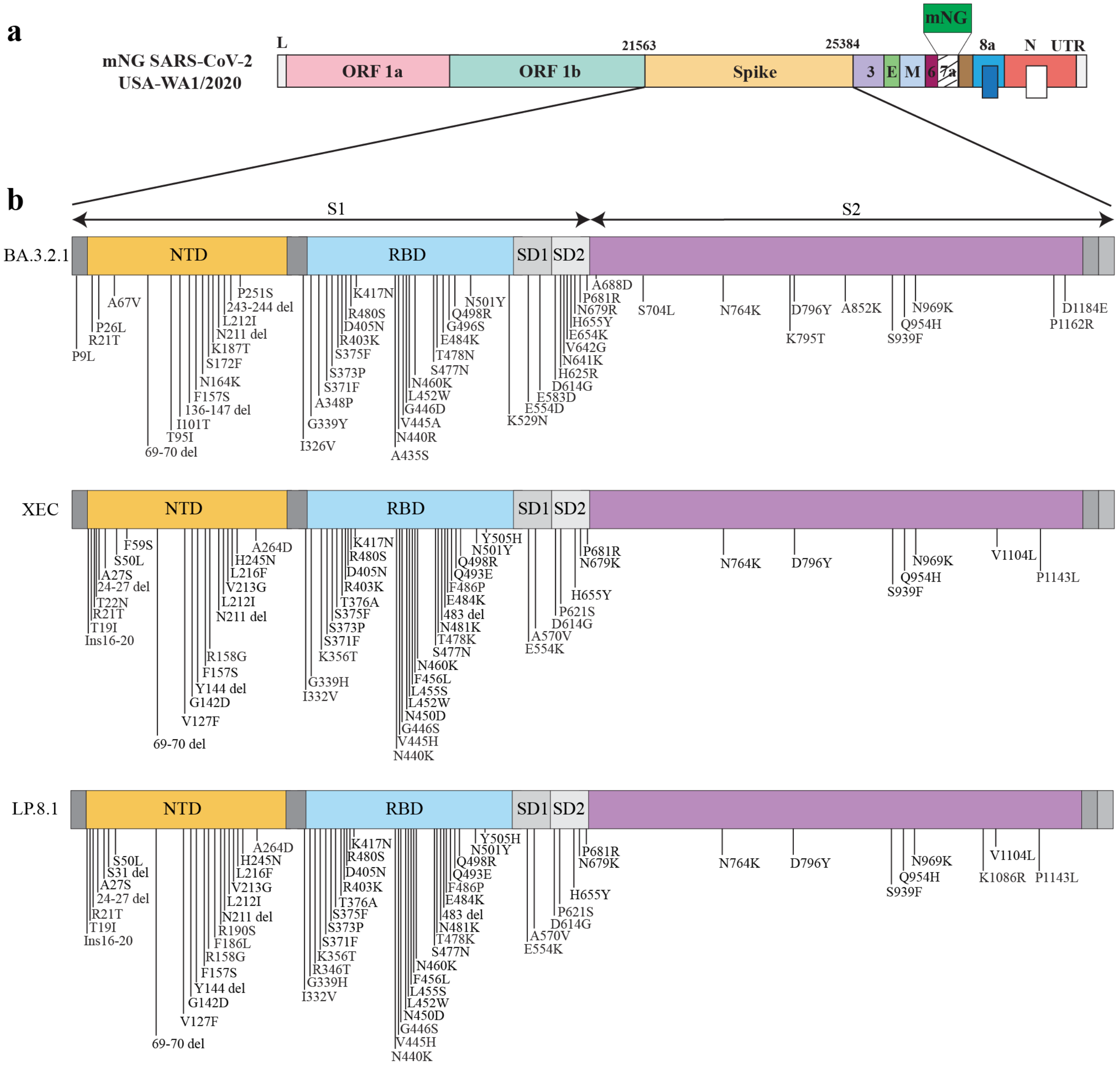


**Extended Data Fig. 1 Schematic of mNG SARS-CoV-2 spike-variant construction**. **a,** The mNG SARS-CoV-2 derived from strain USA-WA1/2020 (WA1) was used as the backbone. The WA1 spike ORF was replaced with the spike gene from each variant to generate the corresponding variant-spike mNG SARS-CoV-2. L, leader sequence; ORF, open reading frame; E, envelope protein; M, membrane protein; N, nucleocapsid; UTR, untranslated region; mNG, mNeonGreen. **b,** Spike mutations across BA.3.2.1, XEC, and LP.8.1. Amino acid substitutions in each spike variant relative to WA1 are annotated along the domain map. NTD: N-terminal domain of spike; RBD: receptor binding domain; S1: N-terminal furin-cleavage fragment of S; SD: subdomain; S2: C-terminal furin-cleavage fragment of S.

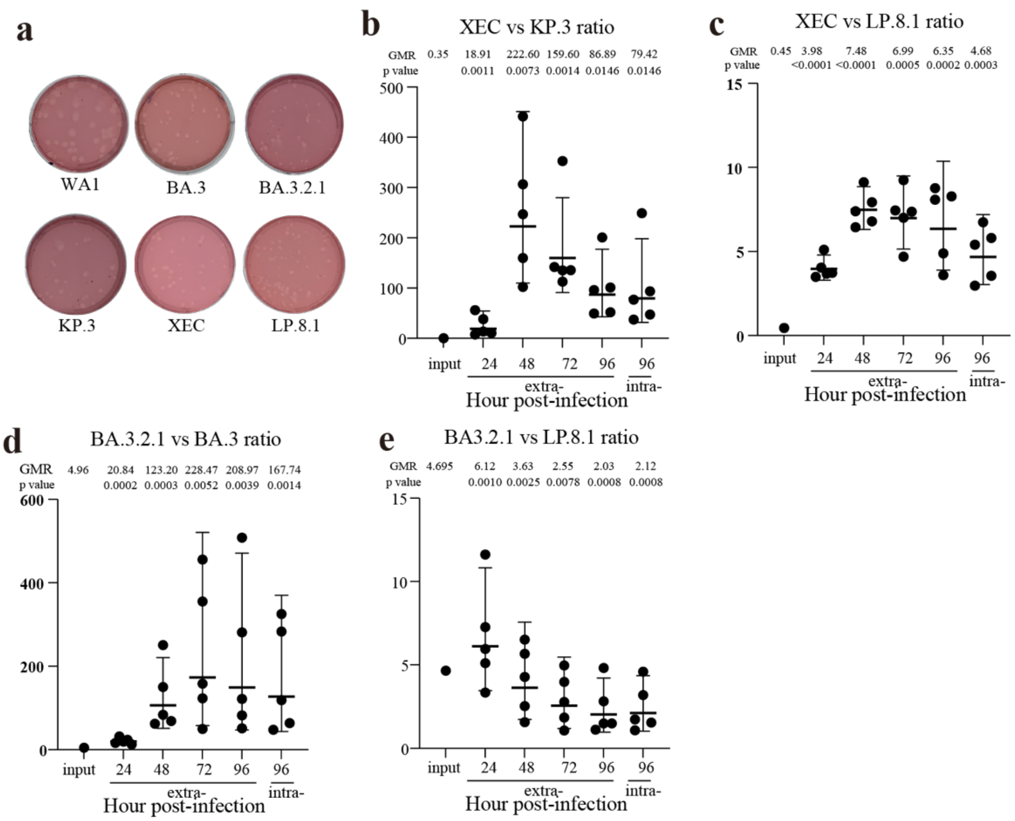


**Extended Data Fig. 2 Characterization of mNG SARS-CoV-2 spike variants**. **a.** Representative plaque morphologies of mNG SARS-CoV-2 variants on Vero E6-TMPRSS2 cells. **b-e.** Scatter plot of RNA ratios measured in infected HAE: XEC: KP.3 **(b)**, XEC: LP.8.1 **(c)**, BA.3.2.1: BA.3 **(d)**, and BA.3.2.1: LP.8.1**(e)**. Geometric mean ratios (GMRs) and 95% confidence intervals (CIs) are shown. P values were calculated from two-sided linear regression analyses comparing the observed RNA ratio at each time point to the input RNA ratio.

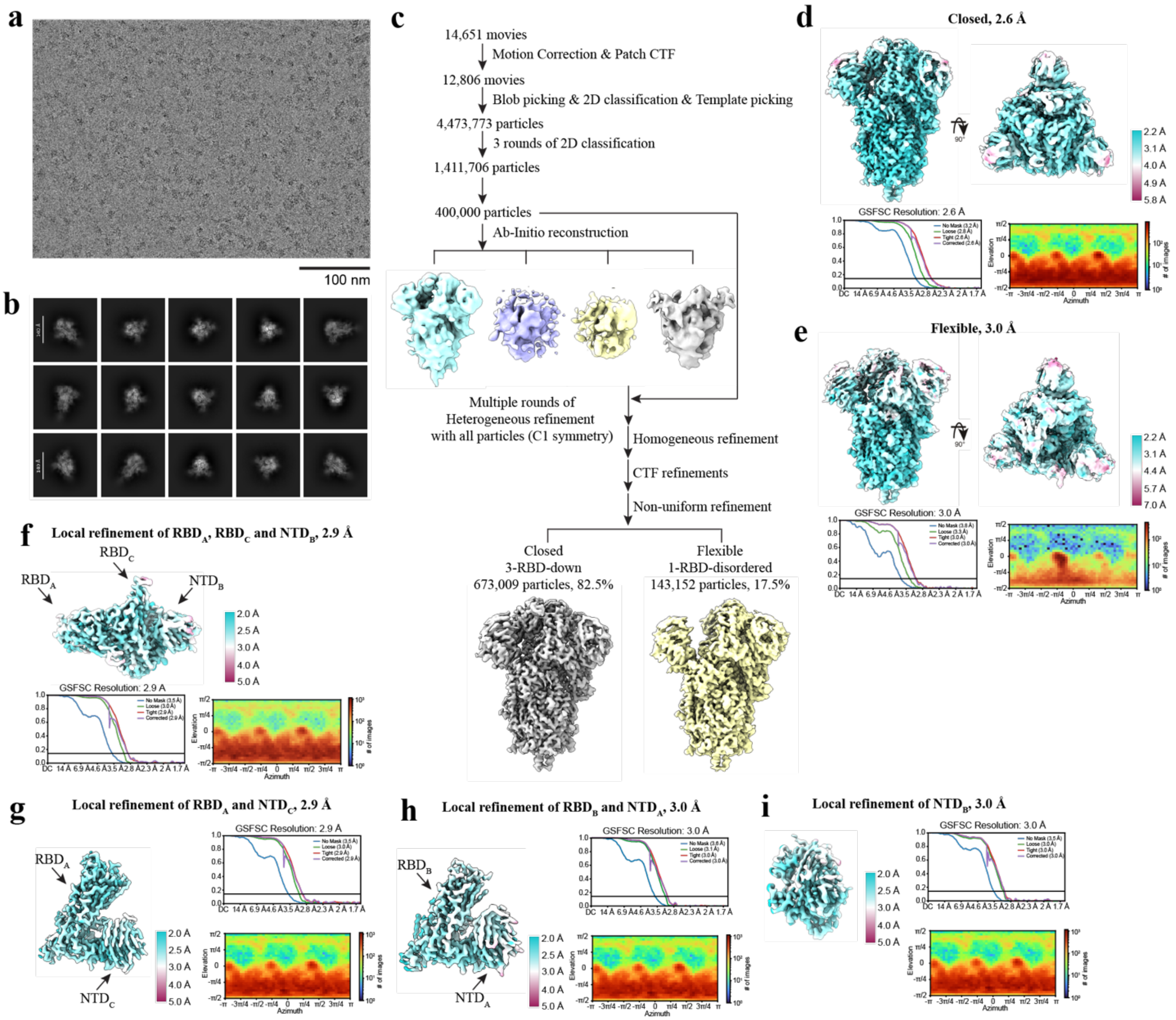


**Extended Data Fig. 3 Cryo-EM processing workflow for the SARS-CoV-2 BA.3.2.1 spike. a,** Representative micrograph of the BA.3.2.1 spike. Scale bar, 100 nm. **b.** Representative 2D class averages of selected particles. **c,** Data-processing flowchart. **d-e,** Global map quality for the non-uniform refinement maps of the closed (**d**) and flexible (**e**) conformations, showing local-resolution distributions, Fourier shell correlation (FSC) curves, and particle angular distributions. **f-i,** Local refinement of the $\text{RBD}_A$, $\text{RBD}_C$, and $\text{NTD}_B$ **(f)**, $\text{RBD}_A$ and $\text{NTD}_C$ **(g)**, $\text{RBD}_B$ and $\text{NTD}_A$ **(h)**, and $\text{NTD}_B$ **(i)** in closed conformation, with corresponding local-resolution distributions, FSC curves, and angular distributions.

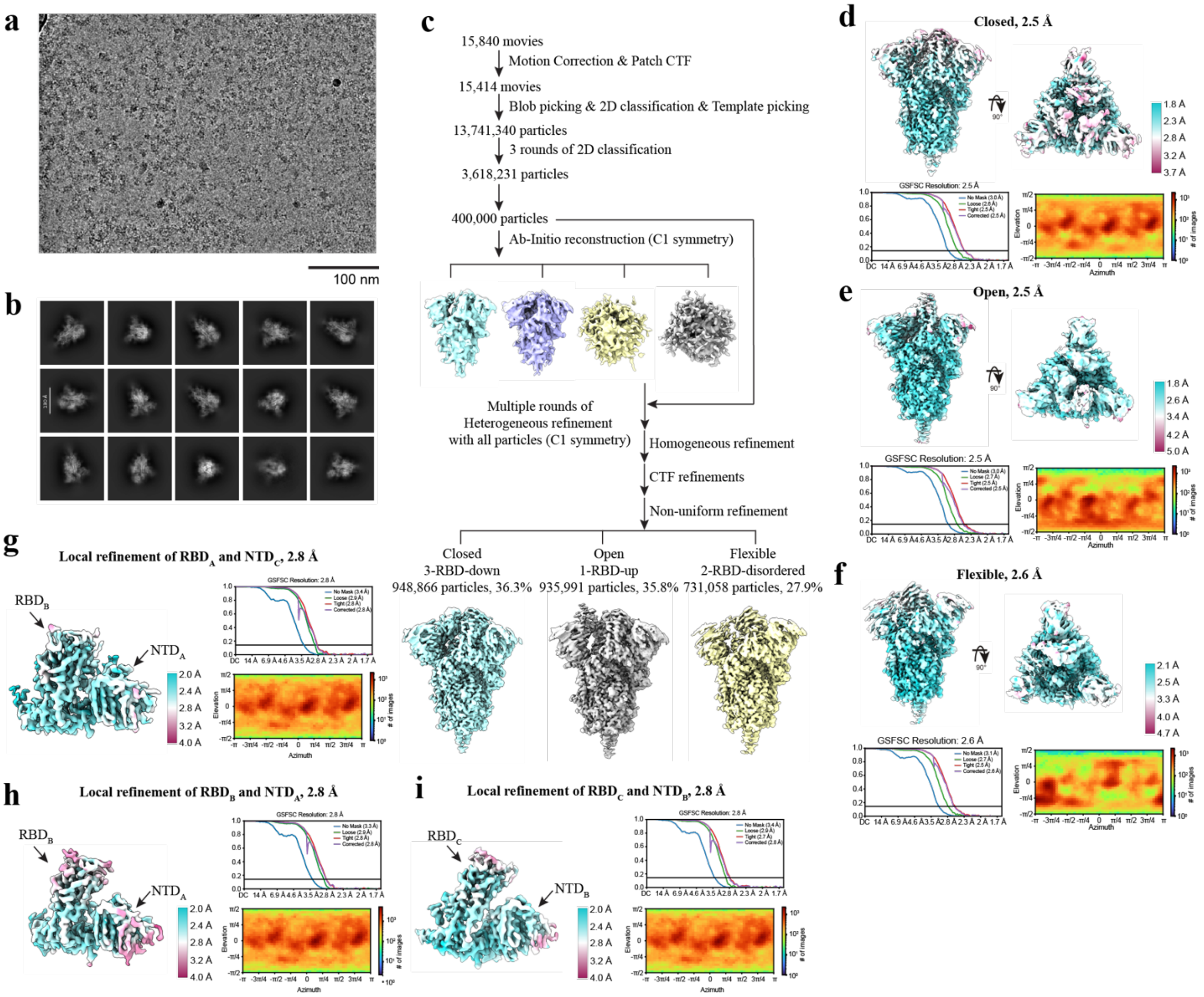


**Extended Data Fig. 4 Cryo-EM processing workflow for the SARS-CoV-2 LP.8.1 spike. a.** Representative micrograph of the LP.8.1 spike. Scale bar, 100 nm. **b.** Representative 2D class averages of selected particles. **c.** Data-processing flowchart. **d-f.** Global map quality for the non-uniform refinement maps of LP.8.1 in the closed (**d**), open (**e**), and flexible (**f**) conformations, showing local-resolution distributions, FSC curves, and particle angular distributions. **g-i,** Local refinements for the closed conformation, with corresponding local-resolution distributions, FSC curves, and angular distributions for: RBD$_A$ and NTD$_C$ **(g)**, RBD$_B$ and NTD$_A$ **(h)**, RBD$_C$ and NTD$_B$ **(i)**.

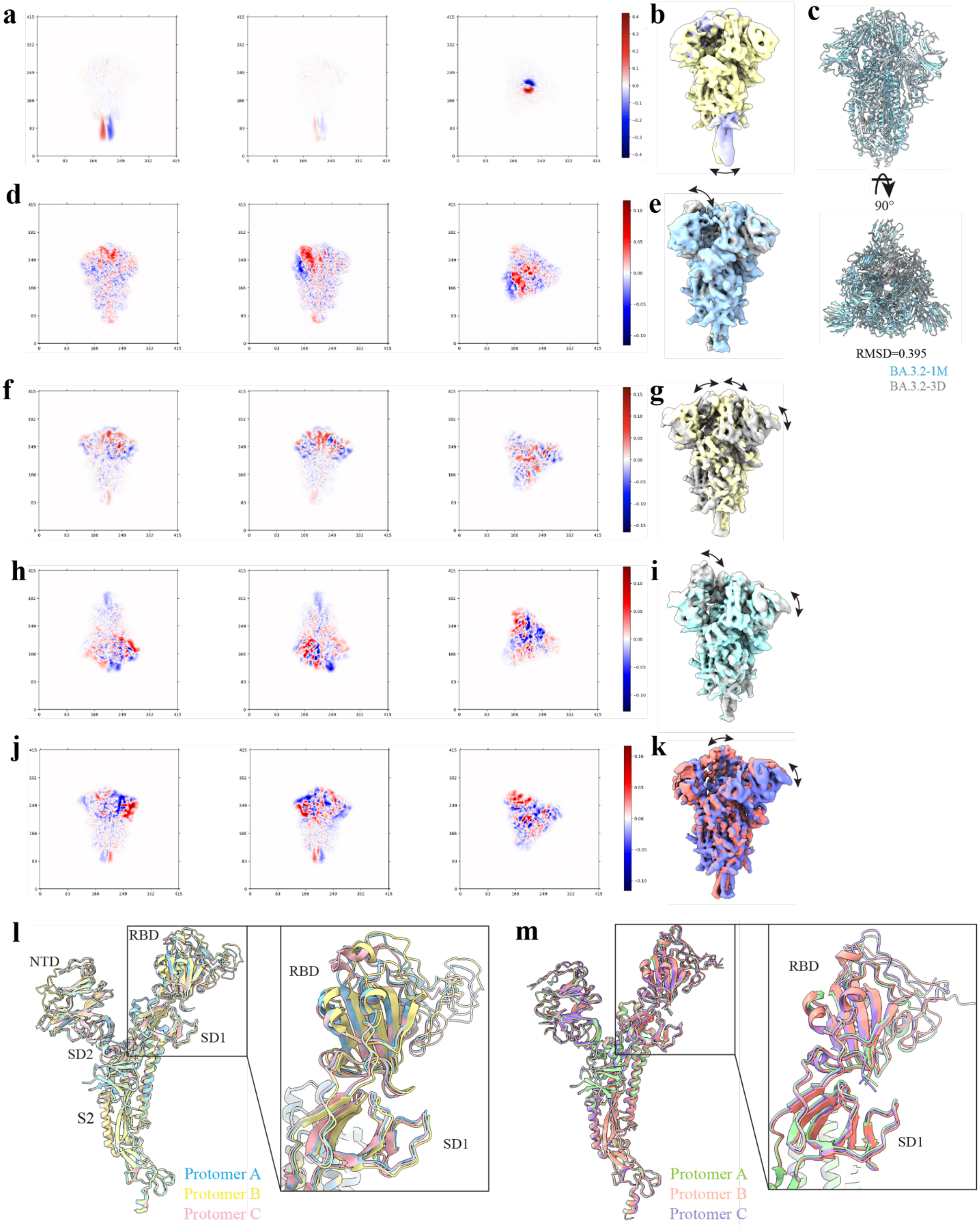


**Extended Data Fig. 5 Structural analysis of BA.3.2.1 and LP.8.1 spike. a,** Scatter plots showing the dispersion of particle latent coordinates throughout the closed BA.3.2.1 spike ensemble. **b,** Representative cryo-EM maps of the closed BA.3.2.1 spike after 3D flexible-refinement training. **c,** Structural alignment of the BA.3.2.1 spike in the closed (gray) and flexible (blue) conformations. **d,** Scatter plots showing the dispersion of particle latent coordinates throughout the flexible BA.3.2.1 spike ensemble. **e,** Representative cryo-EM maps of the flexible BA.3.2.1 spike after 3D flexible-refinement training. **f,** Scatter plots showing the dispersion of particle latent coordinates throughout the closed LP.8.1 spike ensemble. **g,** Representative cryo-EM maps of the closed LP.8.1 spike after 3D flexible-refinement training. **h,** Scatter plots showing the dispersion of particle latent coordinates throughout the open LP.8.1 spike ensemble. **i.** Representative cryo-EM maps of the open LP.8.1 spike after 3D flexible-refinement training. **j.** Scatter plots showing the dispersion of particle latent coordinates throughout the flexible LP.8.1 spike ensemble. **k,** Representative cryo-EM maps of the flexible LP.8.1 spike after 3D flexible-refinement training. **l-m,** Superposition of the main-chain protomers of the BA.3.2.1 spike (**l**) or LP.8.1 spike (**m**) in the closed conformations, aligned using S2 residues 908-1035 of the HR1-CH region. Right panels: magnified views of the RBD and SD1 regions.

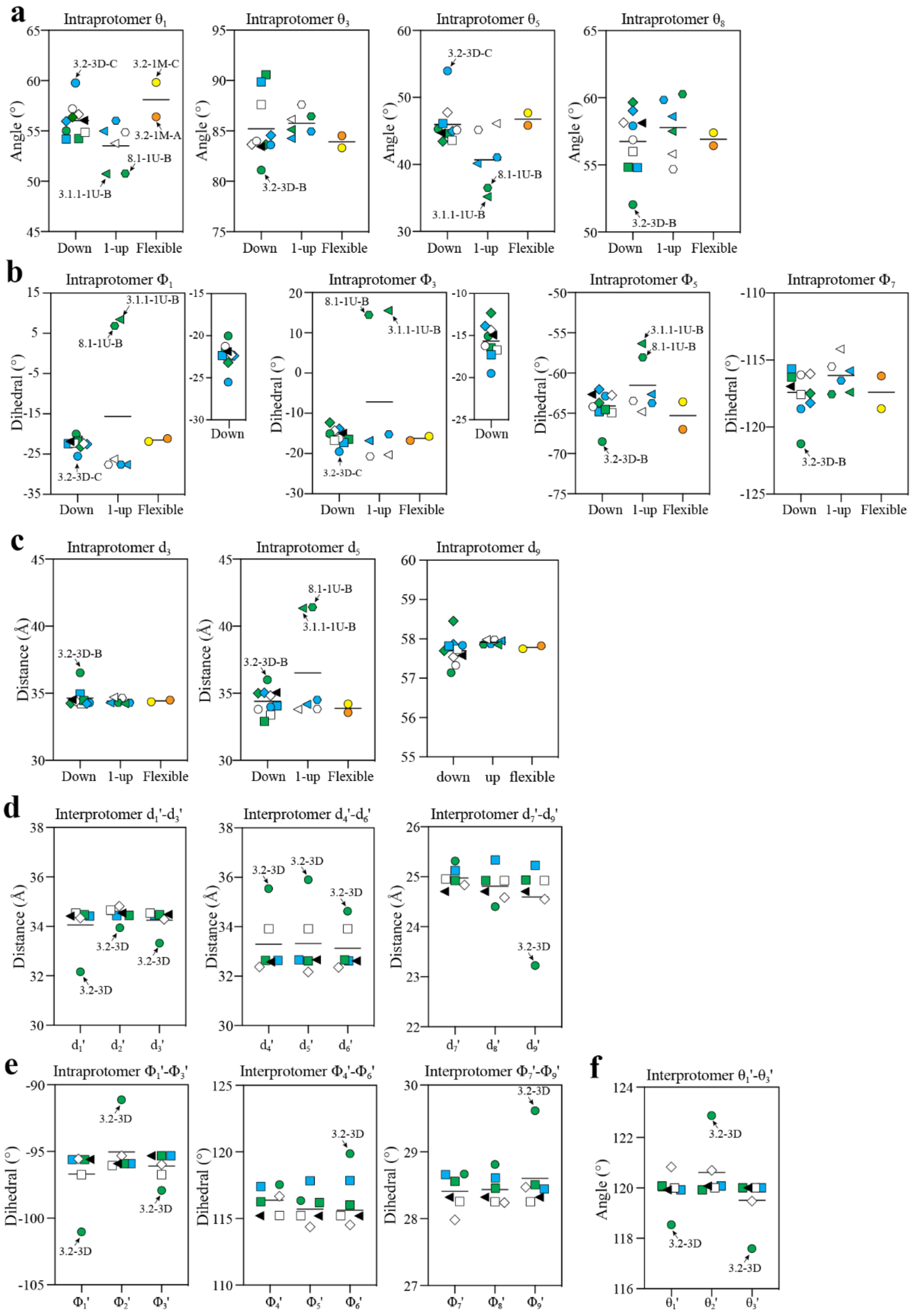


**Extended Data Fig. 6 Vector analysis of the BA.3.2.1 spike. a,** Intraprotomer angles $\theta_1$, $\theta_3$, $\theta_5$, and $\theta_8$ describing angular motion of SD2 relative to RBD about SD1, NTD' relative to SD2 about SD1, SD1 relative to RBD about RBDa, and CD relative to SD2 about SD1, respectively. **b,** Intraprotomer $\Phi_1$, $\Phi_3$, $\Phi_5$, $\Phi_7$ dihedral angles describing rotation of the SD2 and RBDa about SD1 to RBD axis, NTD' and RBD about SD2 to SD1 axis, the NTD and SD1 about SD2 to NTD′ axis, and the SD1 and S2s about SD2 to CD axis, respectively, within the same protomer. **c,** Intraprotomer $d_3$, $d_5$, and $d_9$ describing distances between SD2-

SD1, SD1-RBD, and CD-SD2, respectively, within the same protomer. **d.** Interprotomer $d_1'$-$d_3'$, $d_4'$-$d_6'$, and $d_7'$-$d_9'$ describing the distances between RBDs, RBD-NTD, and SD1-NTD measured across protomer pairs A-C, C-B, and B-A, respectively. **e.** Interprotomer $\Phi_1'$-$\Phi_3'$, $\Phi_4'$-$\Phi_6'$, and $\Phi_7'$-$\Phi_9'$ dihedral angles describing rotation between two RBDa from neighboring protomers about the RBD-RBD axis, SD1a-NTD'a from neighboring protomers about the SD1-NTD' axis, and SD2a from neighboring protomers about the SD1-NTD' axis, respectively. **f.** Interprotomer $\theta_1'$-$\theta_3'$ angles formed by $RBD_A$-$RBD_C$-$RBD_B$ ($\theta_1'$), $RBD_C$-$RBD_B$-$RBD_A$ ($\theta_2'$), and $RBD_b$-$RBD_a$-$RBD_c$ ($\theta_3'$).

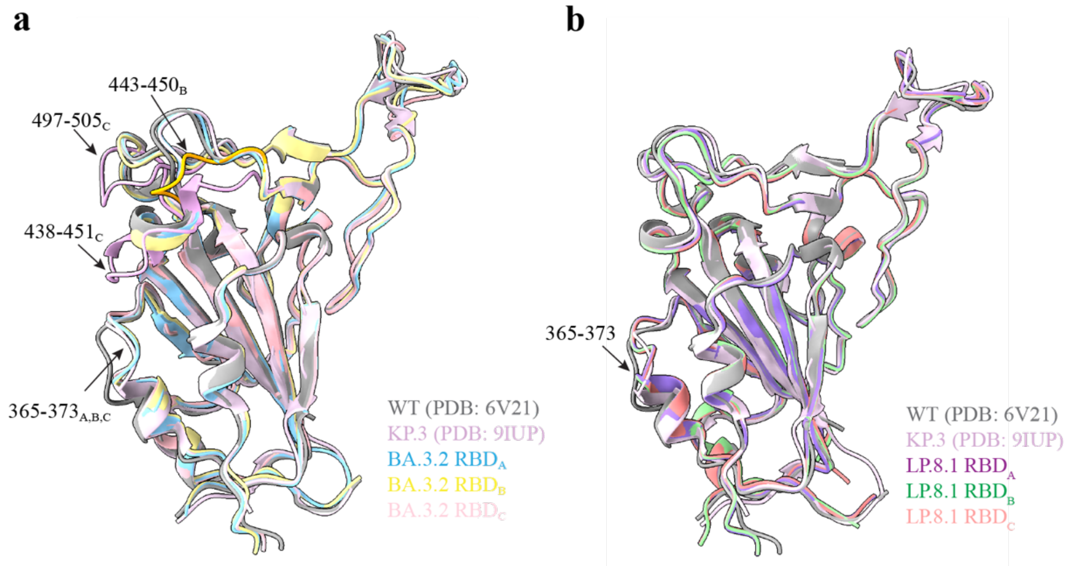


**Extended Data Fig. 7 Structural rearrangements of the BA.3.2.1 RBD. a.** Structural alignment of the BA.3.2.1 (closed; 3D) RBDs from the three protomers with WA1 RBD and KP.3 RBD. The 445–450 loop in protomer B of closed BA.3.2.1 is indicated by an arrow and colored orange; the 438–451 and 497–505 loops in protomer C of closed BA.3.2.1 are indicated by arrows and colored plum. **b.** Structural alignment of the LP.8.1 (closed; 3D) RBDs from the three protomers with WA1 RBD and KP.3 RBD. The loop 365-273 in (**a-b**) is annotated.

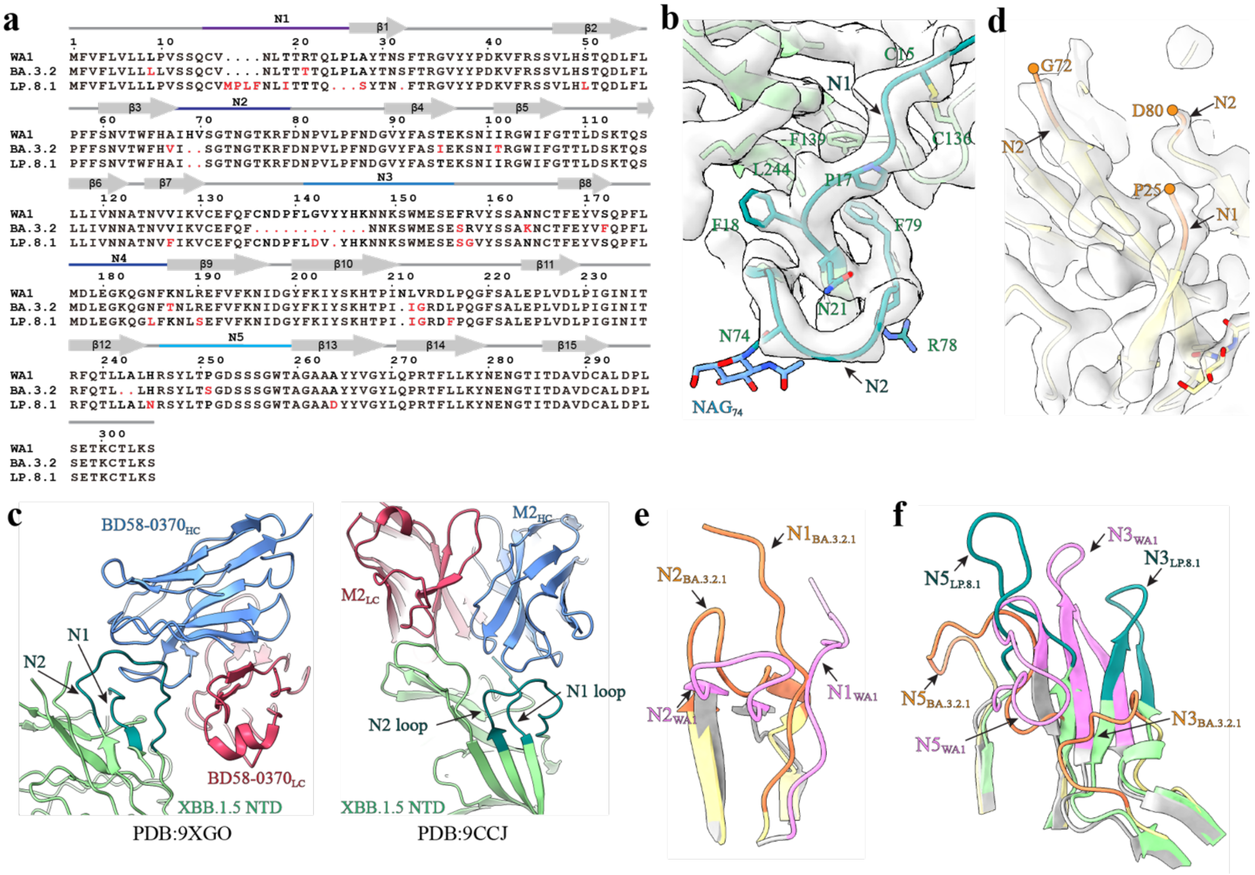


**Extended Data Fig. 8 Structural rearrangements of the BA.3.2.1 NTD. a,** Secondary-structure maps and sequence alignment of WA1, BA.3.2.1, and LP.8.1 NTDs. Mutations are highlighted in red. **b,** Magnified view of the interaction network stabilizing the thread-through arrangement of N1 and N2 loops in the LP.8.1 NTD, with the cryo-EM density shown in gray. **c.** N1-N2 loop crossing in XBB.1.5 NTD stabilized by NAbs BD58-0370 (left) or M2 (right). **d.** Magnified view of the N1-N2 loop region in the BA.3.2.1 NTD, with the cryo-EM density shown in gray. **e.** Zoomed-in superposition of the N1 and N2 loops from WA1 NTD (plum, PDB 76B2) and BA.3.2.1 NTD (orange). **f.** Zoomed-in superposition of the N3 and N5 loops from WA1 NTD (plum), BA.3.2.1 NTD (orange), and LP.8.1 NTD (teal).

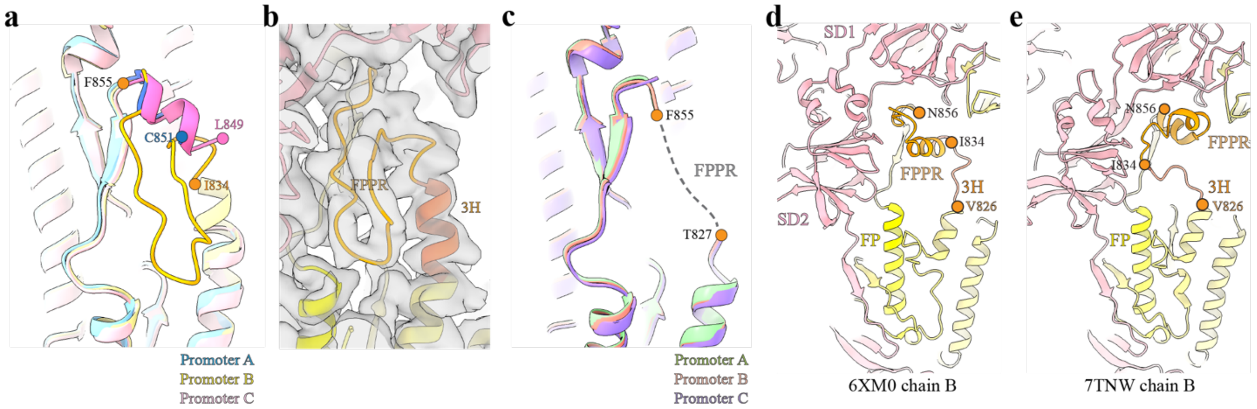


**Extended Data Fig. 9 Structures of spike FPPRs. a,** Alignment of the FPPR across three protomers of closed BA.3.2.1 spike. In protomer B, the FPPR is highlighted in orange. **b,** Magnified view of the FPPR in protomer B of the closed BA.3.2.1 spike, with cryo-EM density shown in gray. **c,** Alignment of the FPPR across the three protomers of the closed LP.8.1 spike. **d-e**, Reference views of the FPPR (orange) and FP (yellow) in WA1 (**d**) and Omicron B.1.1.529 (**e**) spike structures.

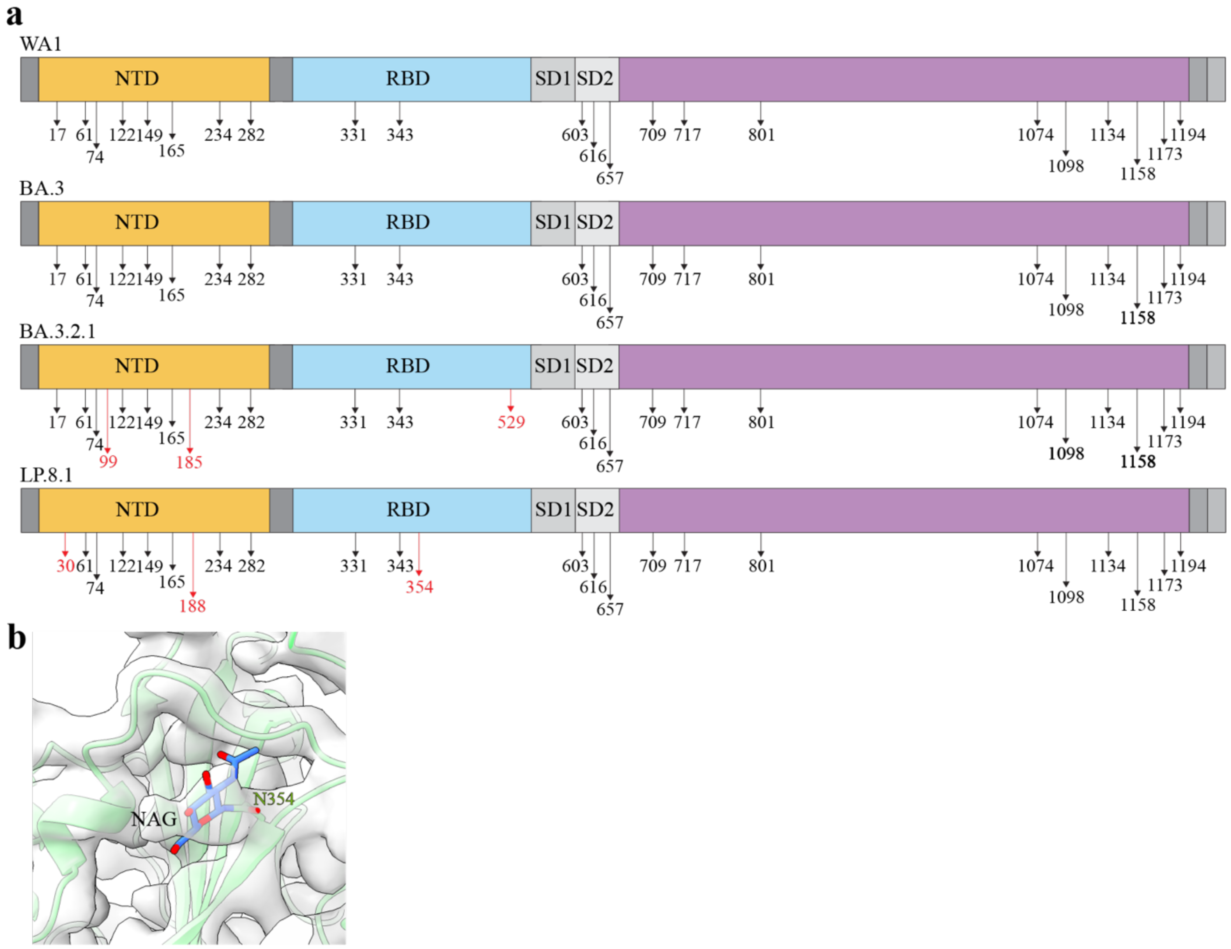


**Extended Data Fig. 10 N-linked glycans on BA.3.2.1 and LP.8.1 Spikes. a,** Schematic map of *N*-linked glycosylation sites on the BA.3.2.1 and LP.8.1 spikes. Additional glycosylation sites unique to BA.3.2.1 are highlighted in red. **b,** Magnified view of the N354 glycan on the LP.8.1 spike, with the cryo-EM density shown in gray.

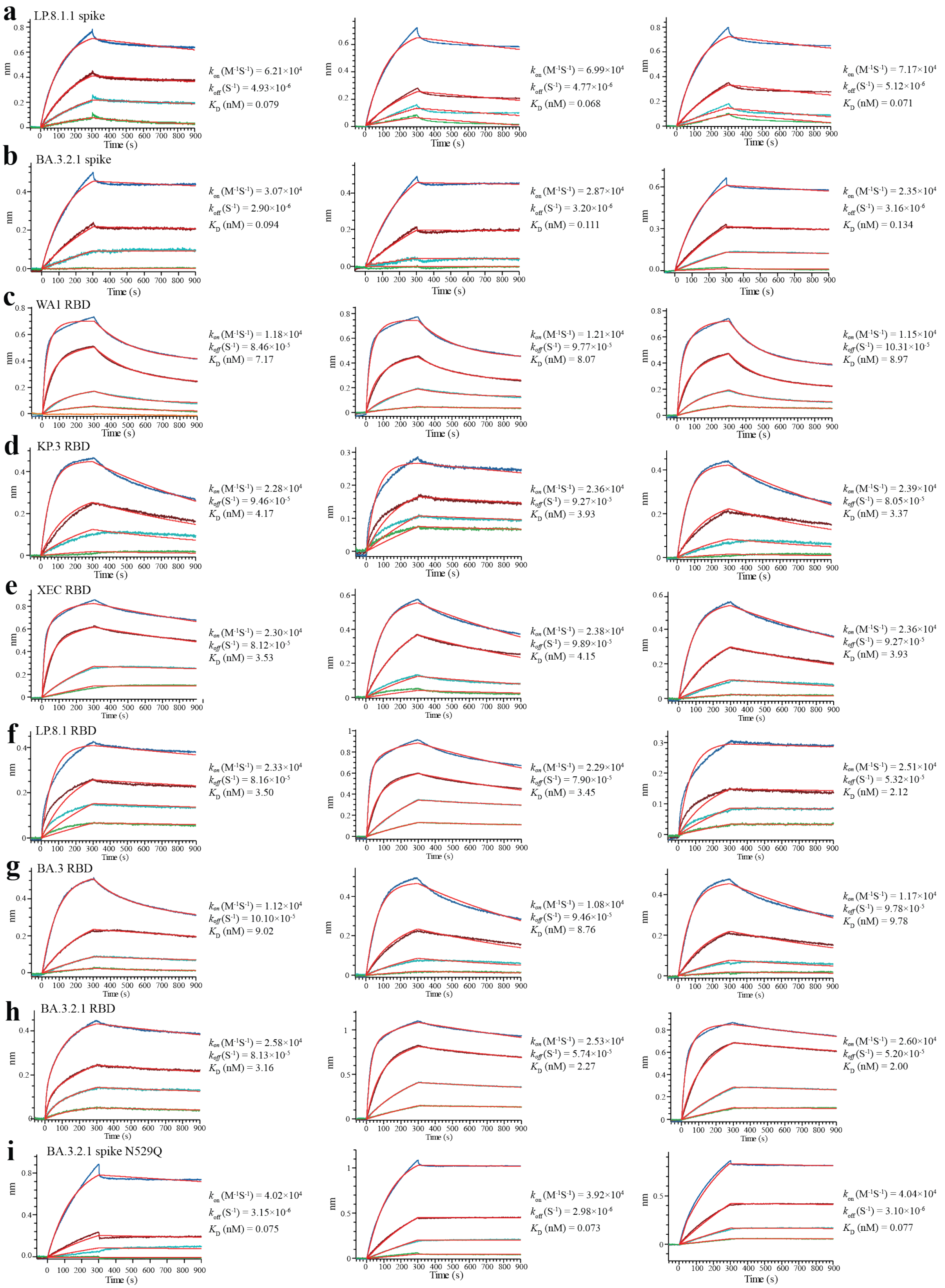


**Extended Data Fig. 11 BLI analysis of hACE2-RBD binding. a-b,** Representative sensorgrams showing binding of trimeric spikes to hACE2: LP.8.1 spike (**a**) and BA.3.2.1 spike (**b**). **c-h**. Sensorgrams for hACE2

binding to RBDs from WA1 (**c**), KP.3 (**d**), XEC (**e**), LP.81 (**f**), BA.3 (**g**), and BA.3.2.1 (**h**). **I,** Sensorgrams for BA.3.2.1 spikes with N529Q mutation binding to hACE2. For each panel, $k_{on}$, $k_{off}$, and $K_D$ from each independent repeat are listed to the right. Thin colored traces represent raw association/dissociation signals; red curves indicate the global fit to the binding model. All measurements were performed in triplicate under identical buffer and temperature conditions.

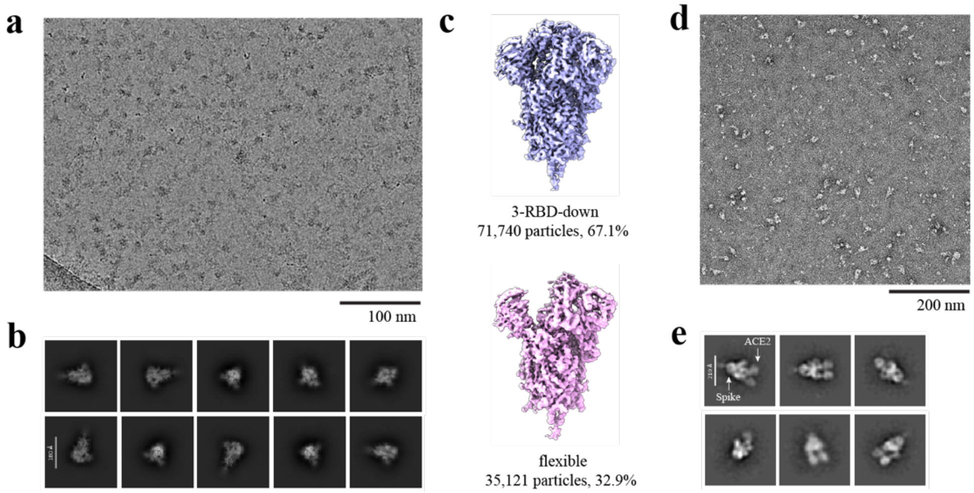


**Extended Data Fig. 12 BA.3.2.1 spike/hACE2 complexes at different temperatures. a,** Representative cryo-EM micrograph of the BA.3.2.1 spike/hACE2 sample incubated at 4°C. Scale bar, 100 nm. **b,** Representative 2D class averages of selected particles in (**a**). **c,** Cryo-EM reconstructions g from the dataset in (**a**), showing only apo BA.3.2.1 spike in the closed (left) and flexible (right) conformations only (no hACE2 bound). **d,** Representative negative-stain EM micrograph of the BA.3.2.1 spike/hACE2 sample incubated at 37°C. Scale bar, 200 nm. **e,** Representative 2D class averages of selected particles from the 37°C dataset in (d).

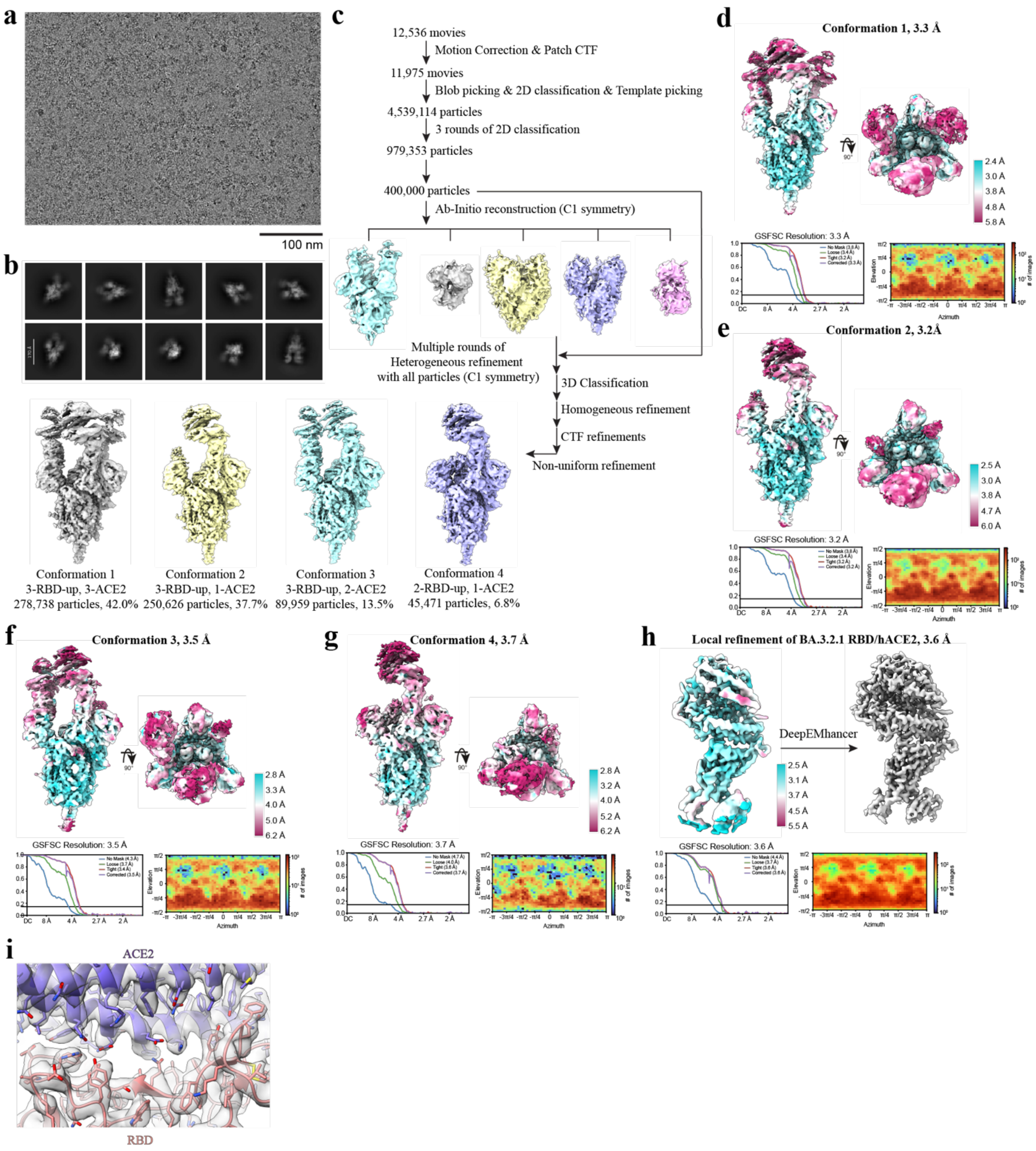


**Extended Data Fig. 13 Cryo-EM processing workflow for the BA.3.2.1 spike-hACE2 complex. a,** Representative micrograph of the BA.3.2.1 spike-hACE2 complex (37°C incubation). Scale bar, 100 nm. **b,** Representative 2D class averages of selected particles. **c,** Data-processing flowchart. **d-g,** Global map quality for the non-uniform refinement maps of the BA.3.2.1 spike–hACE2 complex in conformation 1 (**d**), conformation 2 (**e**), conformation 3 (**f**), and conformation 4 (**g**), showing local-resolution distributions, FSC curves, and particle angular distributions. **h.** Local refinement of the RBD-hACE2 region for conformation 1, with corresponding local-resolution distribution, FSC curve, and angular distribution of the locally refined map. **i.** Overall cryo-EM density at the BA.3.2.1 RBD-hACE2 binding interface. Interface residues are shown as sticks with the corresponding cryo-EM density rendered for clarity.

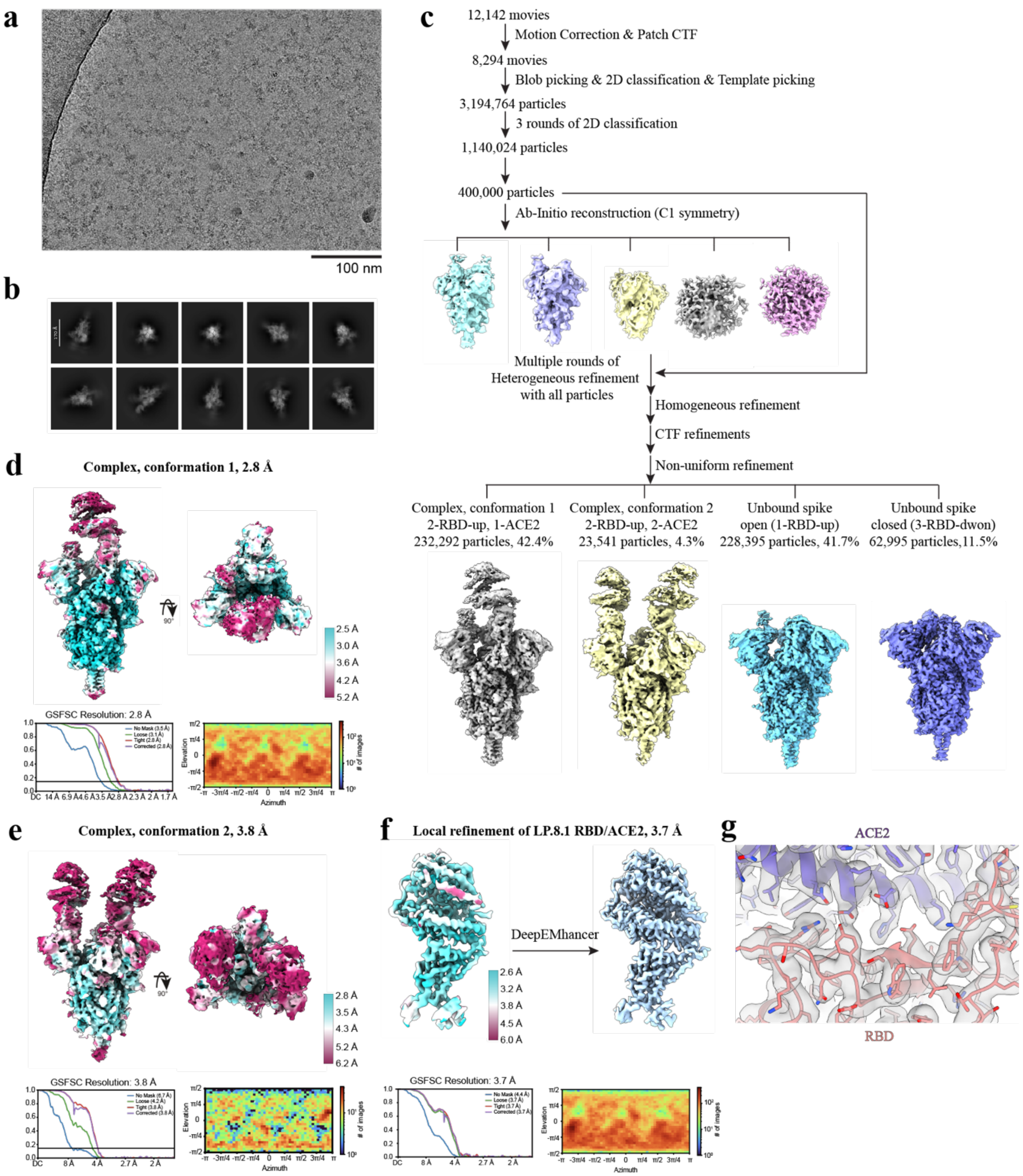


**Extended Data Fig. 14 Cryo-EM processing workflow for the LP.8.1 spike-hACE2 complexes. a,** Representative micrograph of the LP.8.1 spike-hACE2 complex (ice incubation). Scale bar, 100 nm. **b,** Representative 2D class averages of selected particles. **c,** Data-processing flowchart. **d-e,** Global map quality for the LP.8.1 spike-hACE2 complex in conformation 1 (**d**) and conformation 2 (**e**), showing local-resolution distributions, FSC curves, and particle angular distributions. **f,** Local refinement of the LP.8.1 RBD-hACE2 region for conformation 1, with corresponding local-resolution distribution, FSC curve, and angular distribution for the locally refined map. **g.** Overall cryo-EM density at the LP.8.1 RBD-hACE2 binding interface. Interface residues are shown as sticks with the corresponding cryo-EM density rendered for clarity.

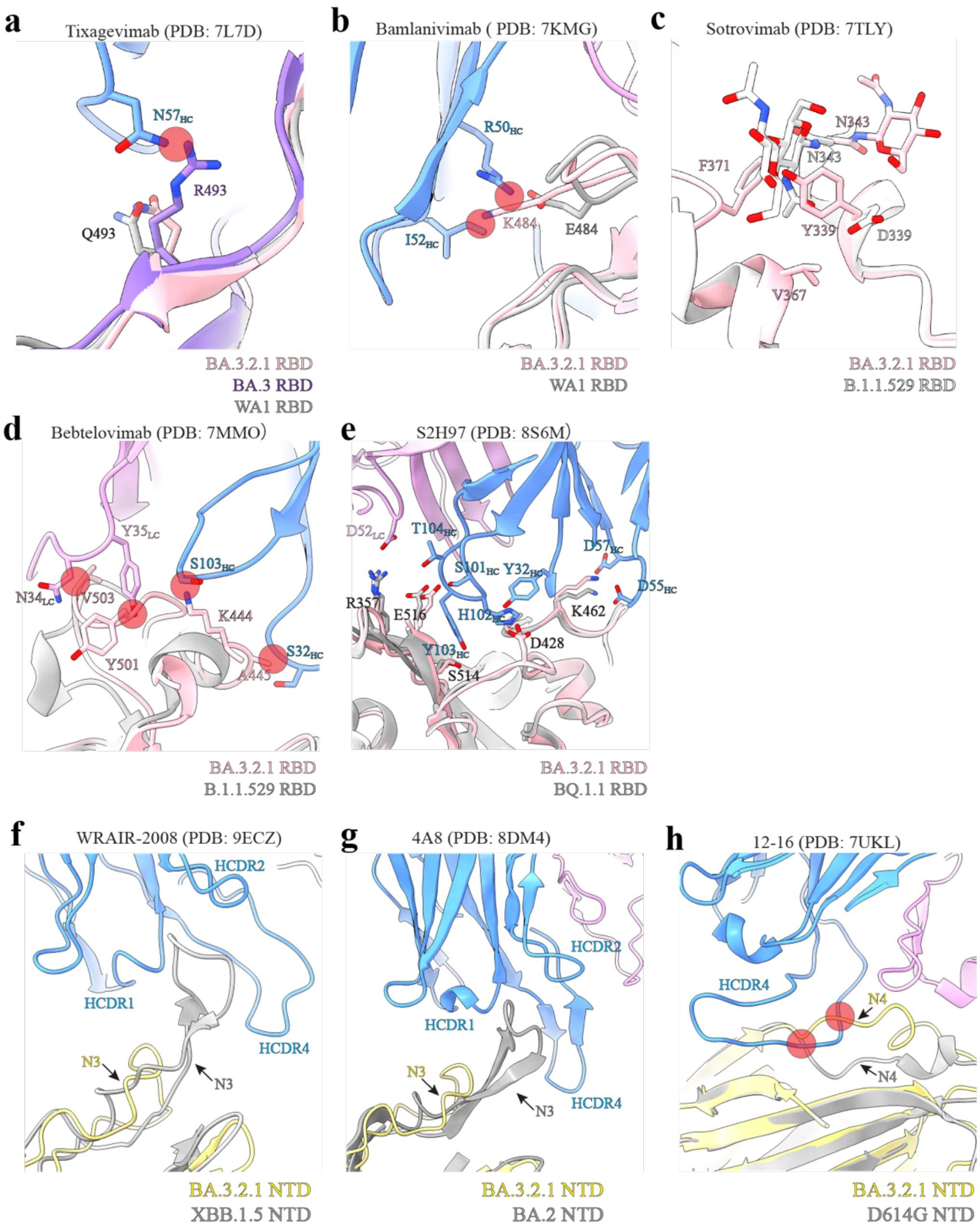


**Extended Data Fig. 15 Structural basis for altered antigenic properties of the BA.3.2.1 spike. a,** Superposition of Tixagevimab/WA1 RBD with BA.3 RBD (PDB 7XB1) and BA.3.2.1 RBD. **b.** Superposition of Bamlanivimab/WA1 RBD with BA.3.2.1 RBD. **c,** Structural alignment of BA.3.2.1 RBD and BA.1.1.529 RBD at the hydrophobic cluster around Y339, V367, and F371, and N343 glycan. **d,** Superposition of Bebtelovimab/B.1.1.529 RBD with BA.3.2.1 RBD. **e,** Superposition of S2H97/WA1 RBD with BA.3.2.1 RBD. **f,** Superposition of XBB.1.5 NTD/WRAIR-2008 onto the BA.3.2.1 NTD. **g,** Superposition of BA.2 NTD/4A8 onto the BA.3.2.1 NTD. **h**, Superposition of D614G NTD/12-16 onto the BA.3.2.1 NTD. In all panels, RBD/NTD residues that interfere with or participate in antibody binding and the corresponding antibody residues are shown as sticks. Steric clashes are indicated by red circles. Antibody heavy and light chains are colored blue and plum, respectively.

**Supplementary Table 1**. Twenty-six human serum samples collected 21-99 days after KP.2-infection

| Serum ID | Age (year) | Gender (F/M) | Race or Ethnicity | Serum collection day (post-KP.2 PCR$^{+}$) | Serum collection date | Doses of parental mRNA vaccine | *FFRNT$_{50}$ | | | | | |
|---|---|---|---|---|---|---|---|---|---|---|---|---|
| | | | | | | | BA.3.2.1-spike | KP.3-spike | XEC-spike | LP.8.1-spike | BA.3-spike | KP.2-spike |
| 1 | 23 | F | Hispanic | 41 | 6/5/24 | 1 dose | 226 | 160 | 101 | 80 | 905 | 160 |
| 2 | 22 | F | Black | 58 | 5/29/24 | 2 doses | 40 | 80 | 100 | 320 | 160 | 80 |
| 3 | 35 | F | Black | 55 | 6/5/24 | 2 doses | 80 | 160 | 100 | 160 | 640 | 160 |
| 4 | 65 | M | Black | 59 | 6/9/24 | 3 doses | 40 | 80 | 101 | 113 | 80 | 80 |
| 5 | 23 | F | Black | 28 | 7/23/24 | none | 10 | 160 | 107 | 453 | 80 | 226 |
| 6 | 42 | F | Hispanic | 24 | 7/29/24 | 2 doses | 640 | 453 | 105 | 640 | 5120 | 1280 |
| 7 | 79 | F | American Indian | 23 | 8/7/24 | 2 doses | 905 | 2560 | 98 | 5120 | 1280 | 3620 |
| 8 | 70 | M | White | 36 | 7/28/24 | none | 40 | 40 | 89 | 80 | 80 | 80 |
| 9 | 30 | F | White | 21 | 8/12/24 | 2 doses | 160 | 160 | 93 | 160 | 640 | 160 |
| 10 | 76 | F | Hispanic | 39 | 7/25/24 | 3 doses | 1810 | 1280 | 93 | 905 | 5120 | 1280 |
| 11 | 80 | F | White | 42 | 7/25/24 | 4 doses | 226 | 320 | 87 | 640 | 640 | 320 |
| 12 | 29 | F | White | 66 | 7/1/24 | none | 40 | 160 | 84 | 113 | 640 | 320 |
| 13 | 82 | F | White | 27 | 8/10/24 | none | 20 | 320 | 84 | 320 | 113 | 320 |
| 14 | 22 | F | Hispanic | 43 | 7/24/24 | none | 40 | 80 | 84 | 80 | 160 | 40 |
| 15 | 75 | F | Hispanic | 54 | 7/17/24 | 4 doses | 640 | 1280 | 91 | 1280 | 3620 | 1280 |
| 16 | 55 | F | White | 83 | 6/19/24 | 2 doses | 113 | 40 | 85 | 80 | 640 | 113 |
| 17 | 74 | F | White | 27 | 8/15/24 | 2 doses | 113 | 320 | 91 | 320 | 640 | 320 |
| 18 | 53 | F | White | 48 | 7/26/24 | none | 57 | 40 | 88 | 40 | 160 | 40 |
| 19 | 34 | F | Hispanic | 36 | 8/12/24 | none | 80 | 80 | 97 | 160 | 640 | 160 |
| 20 | 34 | F | Hispanic | 52 | 8/10/24 | 2 doses | 40 | 80 | 106 | 80 | 320 | 80 |
| 21 | 29 | M | Black | 32 | 8/30/24 | 2 doses | 80 | 80 | 111 | 28 | 640 | 113 |
| 22 | 62 | M | Hispanic | 99 | 6/28/24 | 4 doses | 113 | 320 | 137 | 160 | 453 | 160 |
| 23 | 26 | F | Hispanic | 73 | 7/25/24 | 3 doses | 160 | 320 | 105 | 640 | 640 | 320 |
| 24 | 28 | F | Asian | 41 | 8/28/24 | 4 doses | 40 | 20 | 84 | 28 | 453 | 40 |
| 25 | 46 | M | Hispanic | 62 | 9/2/24 | 2 doses | 14 | 113 | 98 | 80 | 320 | 226 |
| 26 | 28 | M | Hispanic | 62 | 9/4/24 | none | 57 | 40 | 113 | 20 | 640 | 28 |
| Median | 38.5 | - | - | 42.5 | - | - | - | - | - | - | - | - |
| #GMT | - | - | - | - | - | - | 91 | 156 | 104 | 178 | 484 | 188 |
| †95% CI | - | - | - | - | - | - | 55-153 | 97-251 | 91-101 | 128-292 | 412-741 | 166-340 |

*Individual FFRNT$_{50}$ value is the geometric mean of duplicate FFRNT$_{50}$ results.
#Geometric mean neutralizing titers (GMT).
†95% confidence interval (95% CI) for the GMT.

**Supplementary Table 2**. Twenty-one human serum samples collected 23-105 days after KP.3-infection

| Serum ID | Age (year) | Gender (F/M) | Race or Ethnicity | Serum collection day (post-KP.3 PCR$^{+}$) | Serum collection date | Doses of parental mRNA vaccine | *FFRNT$_{50}$ | | | | | |
|---|---|---|---|---|---|---|---|---|---|---|---|---|
| | | | | | | | BA.3.2.1-spike | KP.3-spike | XEC-spike | LP.8.1-spike | BA.3-spike | KP.2-spike |
| 1 | 91 | F | White | 23 | 8/8/24 | 5 doses | 14 | 57 | 99 | 80 | 20 | 57 |
| 2 | 54 | M | White | 48 | 8/15/24 | 2 doses | 160 | 320 | 104 | 320 | 905 | 453 |
| 3 | 67 | M | White | 35 | 8/17/24 | 3 doses | 40 | 40 | 104 | 80 | 320 | 40 |
| 4 | 26 | F | Black | 33 | 8/22/24 | none | 28 | 160 | 106 | 226 | 160 | 160 |
| 5 | 34 | F | Black | 33 | 8/22/24 | none | 80 | 320 | 106 | 320 | 1280 | 640 |
| 6 | 68 | F | Black | 36 | 8/25/24 | 6 doses | 80 | 80 | 102 | 160 | 640 | 113 |
| 7 | 29 | F | White | 34 | 8/28/24 | 2 doses | 320 | 905 | 103 | 1280 | 2560 | 1280 |
| 8 | 58 | F | Hispanic | 79 | 9/7/24 | 3 doses | 20 | 40 | 84 | 40 | 113 | 28 |
| 9 | 59 | F | Black | 53 | 9/9/24 | 6 doses | 320 | 80 | 89 | 80 | 1280 | 80 |
| 10 | 59 | M | White | 50 | 9/9/24 | 5 doses | 80 | 80 | 82 | 80 | 640 | 113 |
| 11 | 48 | F | Hispanic | 40 | 9/16/24 | 2 doses | 160 | 160 | 95 | 320 | 905 | 320 |
| 12 | 62 | F | White | 56 | 9/17/24 | 3 doses | 80 | 640 | 100 | 453 | 5120 | 453 |
| 13 | 37 | M | Hispanic | 54 | 9/20/24 | 3 doses | 40 | 40 | 98 | 40 | 320 | 28 |
| 14 | 27 | F | Black | 53 | 10/3/24 | none | 40 | 113 | 110 | 160 | 320 | 80 |
| 15 | 45 | F | White | 39 | 10/3/24 | 2 doses | 113 | 160 | 131 | 160 | 226 | 160 |
| 16 | 71 | M | White | 72 | 10/5/24 | none | 10 | 1280 | 135 | 5120 | 1280 | 1280 |
| 17 | 53 | M | White | 27 | 10/8/24 | none | 40 | 320 | 91 | 320 | 1280 | 160 |
| 18 | 82 | F | White | 87 | 10/31/24 | 6 doses | 160 | 640 | 107 | 640 | 1280 | 453 |
| 19 | 96 | F | Black | 105 | 11/8/24 | 3 doses | 80 | 320 | 147 | 453 | 640 | 453 |
| 20 | 48 | F | White | 87 | 12/2/24 | 2 doses | 10 | 80 | 226 | 57 | 40 | 40 |
| 21 | 52 | M | Hispanic | 81 | 12/13/24 | 2 doses | 226 | 1280 | 1280 | 1280 | 640 | 1280 |
| Median | 54 | - | - | 50 | - | - | - | - | - | - | - | - |
| #GMT | - | - | - | - | - | - | 63 | 186 | 108 | 230 | 508 | 186 |
| †95% CI | - | - | - | - | - | - | 51-108 | 134-264 | 97-150 | 180-365 | 364-764 | 171-326 |

*Individual FFRNT$_{50}$ value is the geometric mean of duplicate FFRNT$_{50}$ results.
#Geometric mean neutralizing titers (GMT).
†95% confidence interval (95% CI) for the GMT.

**Supplementary Table 3**. P values for group comparison of GMTs in **Fig. 1i-j**.

| Figure Panels | P values (two-tailed) from the Wilcoxon matched-pairs signed-rank test |
|---|---|
| Fig. 1i | KP.2- versus KP.3-, XEC-, LP.8.1-, BA.3-, BA.3.2.1-spike: 0.0387, <0.0001, 0.9099, 0.0003, 0.001; KP.3- versus XEC-, LP.8.1-, BA.3-, BA.3.2.1-spike: 0.0002, 0.2415, 0.0002, 0.0268; XEC- versus LP.8.1-, BA.3-, BA.3.2.1-spike: 0.0002, <0.0001, 0.2673; LP.8.1- versus BA.3-, BA.3.2.1-spike: 0.0021, 0.0063; BA.3- versus BA.3.2.1: <0.0001. |
| Fig. 1j | KP.3- versus KP.2-, XEC-, LP.8.1-, BA.3-, BA.3.2.1-spike: 0.6819, 0.0005, 0.049, 0.0006, 0.001; KP.2- versus XEC-, LP.8.1-, BA.3-, BA.3.2.1-spike: 0.0012, 0.1310, 0.0007, 0.0002; XEC- versus LP.8.1-, BA.3-, BA.3.2.1-spike: <0.0001, 0.0003, 0.1197; LP.8.1- versus BA.3- BA.3.2.1-spike: 0.0078, 0.0002; BA.3- versus BA.3.2.1 spike: <0.0001. |

**Supplementary Table 4.** List of primers used in this study.

| Primer Name | Sequence (5'-3') | Notes |
|---|---|---|
| CoV-21115V | CATTTGTGGGTTTATACAACAAAAG | For RT-PCR |
| CoV-YH5 | AGCATCCTTGATTTCACC | For RT-PCR |
| K529N-F | CCACTTATGGTGTTGGTTACCAACC | For competition assay of BA.3 and BA.3.2.1 |
| K529N-R | GCAATGTCTCTGCCAAATTGTTGG | For competition assay of BA.3 and BA.3.2.1 |
| A852K-F | CAACAAAGTGACACTTGCAGATGC | For competition assay of BA.3.2.1 and LP.8.1 |
| A852K-R | GTGATTGTACCCGCTAACAGTGC | For competition assay of BA.3.2.1 and LP.8.1 |
| K1086-F | AAGGGCTATCATCTTATGTCCTTCC | For competition assay of LP.8.1 and XEC |
| K1086-R | CAAATGTGTTGTCTGTAGTAATGATTTGTGG | For competition assay of LP.8.1 and XEC |
| F59S-F | CTTTCACACGTGGTGTTTATTACCC | For competition assay of KP.3 and XEC |
| F59S-R | CTTCTCAGTGGAAGCAAAATAAACACC | For competition assay of KP.3 and XEC |
| N529Q-F | GTGGACCTAAACAGTCTACTAATTTGG | For generation BA.3.2.1 spike mutation K529Q |
| N529Q-R | AGTAGACTGTTTAGGTCCACAAACAGTTG | For generation BA.3.2.1 spike mutation K529Q |
| 2019-nCoV_N2-F | TTACAAACATTGGCCGCAAA | For RT-qPCR |
| 2019-nCoV_N2-R | GCGCGACAT TCCGAAGAA | For RT-qPCR |
| β-actin-F | AGAGCTACGAGCTGCCTGAC | For RT-qPCR |
| β-actin-R | AGCACTGTGTTGGCGTACAG | For RT-qPCR |

**Supplementary Table 5.** Statistics for 3D reconstruction and model refinement for BA.3.2.1 spike and BA.3.2.1 spike-ACE2 complex.

| | BA.3.2.1 spike | | BA.3.2.1 spike-hACE2 | | | |
|---|---|---|---|---|---|---|
| | Closed | Flexible | Class 1 | Class 2 | Class 3 | Class 4 |
| EMD | 73393 | 73395 | 73404 | 73405 | 73408 | 73409 |
| PDB | 9YSJ | 9YX6 | - | - | - | - |
| **Data collection and processing** | | | | | | |
| Microscope | Krios | | Krios | | | |
| Camera | K3 | | K3 | | | |
| Voltage (keV) | 300 | | 300 | | | |
| Defocus range (- μm) | 0.9-2.5 | | 0.9-2.5 | | | |
| Pixel size (Å) | 0.832 | | 0.832 | | | |
| Electron dose ($^{-}$ Å$^{-1}$) | 39.84 | | 39.90 | | | |
| Micrographs (no.) | 12,806 | | 11294 | | | |
| **Refinement** | | | | | | |
| Symmetry imposed | C1 | C1 | C1 | C1 | C1 | C1 |
| Particles (no.) | 673,009 | 143,152 | 278,738 | 250,626 | 89,959 | 45,471 |
| Map resolution (Å) | 2.6 | 3.0 | 3.3 | 3.2 | 3.5 | 3.7 |
| Initial model (PDB code) | 7XIY | 7XIY | - | - | - | - |
| **Model composition** | | | | | | |
| Chains | 3 | 3 | - | - | - | - |
| Atoms | 24,392 | 21,789 | - | - | - | - |
| Residues (Protein) | 3,021 | 2,706 | - | - | - | - |
| Ligands (NAG) | 47 | 41 | - | - | - | - |
| **R.m.s. deviations** | | | | | | |
| Bond lengths (Å) | 0.005 | 0.003 | - | - | - | - |
| Bond angles (°) | 0.569 | 0.573 | - | - | - | - |
| **Model statistics** | | | | | | |
| Clash score | 6.93 | 8.84 | - | - | - | - |
| MolProbity score | 1.66 | 1.73 | - | - | - | - |
| Rotamer outliers (%) | 0.75 | 0.71 | - | - | - | - |
| **Ramachandran plot** | | | | | | |
| Outliers (%) | 0.00 | 0.00 | - | - | - | - |
| Allowed (%) | 4.08 | 3.73 | - | - | - | - |
| Favored (%) | 95.92 | 96.27 | - | - | - | - |

**Supplementary Table 6.** Statistics for 3D reconstruction and model refinement for LP.8.1 spike.

| | LP.8.1 spike | | | LP.8.1 spike-ACE2 | | | |
|---|---|---|---|---|---|---|---|
| | Closed | Open | Flexible | Complex | | Apo Spike | |
| | | | | Class 1 | Class 2 | Open | Closed |
| EMD | 73394 | 73535 | 73396 | 73427 | 73428 | 73430 | 73439 |
| PDB | 9YSK | 9YW0 | - | - | - | - | - |
| **Data collection and processing** | | | | | | | |
| Microscope | Krios | | | Krios | | | |
| Camera | K3 | | | K3 | | | |
| Voltage (keV) | 300 | | | 300 | | | |
| Defocus range (- μm) | 0.9-2.5 | | | 0.9-2.5 | | | |
| Pixel size (Å) | 0.832 | | | 0.832 | | | |
| Electron dose ($^{-}$ Å$^{-1}$) | 39.9 | | | 39.8 | | | |
| Micrographs (no.) | 15,414 | | | 8,294 | | | |
| **Refinement** | | | | | | | |
| Symmetry imposed | C1 | C1 | C1 | C1 | C1 | C1 | C1 |
| Particles (no.) | 948,866 | 935,991 | 731,058 | 232,292 | 23,541 | 228,395 | 62,995 |
| Map resolution (Å) | 2.5 | 2.5 | 2.6 | 2.8 | 3.8 | 2.8 | 3.2 |
| Initial model (PDB code) | 9ELI | 9ELL | - | - | - | - | - |
| **Model composition** | | | | | | | |
| Chains | 3 | 3 | - | - | - | - | - |
| Atoms | 24,010 | 24,827 | - | - | - | - | - |
| Residues (Protein) | 2,989 | 3,083 | - | - | - | - | - |
| Ligands | 41 | 46 | - | - | - | - | - |
| **R.m.s. deviations** | | | | | | | |
| Bond lengths (Å) | 0.003 | 0.002 | - | - | - | - | - |
| Bond angles (°) | 0.517 | 0.528 | - | - | - | - | - |
| **Model statistics** | | | | | | | |
| Clash score | 6.62 | 6.92 | - | - | - | - | - |
| MolProbity score | 1.53 | 1.67 | - | - | - | - | - |
| Rotamer outliers (%) | 0.76 | 0.96 | - | - | - | - | - |
| **Ramachandran plot** | | | | | | | |
| Outliers (%) | 0.00 | 0.00 | - | - | - | - | - |
| Allowed (%) | 2.91 | 4.19 | - | - | - | - | - |
| Favored (%) | 97.09 | 95.81 | - | - | - | - | - |

**Supplementary Table 7.** Intra-protomer vector descriptions

| Distance | Centroids | Description |
|---|---|---|
| $d_1$ | NTD, NTD′ | Term describing the distance between NTD and NTD′ within the same protomer |
| $d_2$ | NTD′, SD2 | Term describing the distance between NTD′ and SD2 within the same protomer |
| $d_3$ | SD2, SD1 | Term describing the distance between SD2 and SD1 within the same protomer |
| $d_4$ | SD2, CD | Term describing the distance between SD2 and CD within the same protomer |
| $d_5$ | SD1, RBD | Term describing the distance between SD1 and RBD within the same protomer |
| $d_6$ | CD, S2s | Term describing the distance between CD and S2s within the same protomer |
| $d_7$ | NTDa, NTD | Term describing the distance between NTDa and NTD within the same protomer |
| $d_8$ | RBD, RBDa | Term describing the distance between RBD and RBDa within the same protomer |
| $d_9$ | CD, SD2 | Term describing the distance between CD and SD2 within the same protomer |
| $d_{10}$ | SD1, SD2, | Term describing the distance between SD1 and SD2 within the same protomer |
| Angle | | |
| $\theta_1$ | SD2, SD1, RBD | Term describing in angular motion of SD2 relative to RBD about SD1 |
| $\theta_2$ | NTDa, NTD, NTD′ | Term describing in angular motion of NTDa relative to NTD about NTD′ |
| $\theta_3$ | NTD′, SD2, SD1 | Term describing in angular motion of NTD′ relative to SD2 about SD1 |
| $\theta_4$ | NTD, NTD′, SD2 | Term describing in angular motion of NTD relative to NTD′ about SD2 |
| $\theta_5$ | SD1, RBD, RBDa | Term describing in angular motion of SD1 relative to RBD about RBDa |
| $\theta_6$ | NTD′, SD2, CD | Term describing in angular motion of NTD′ relative to SD2 about CD |
| $\theta_7$ | SD2, CD, S2s | Term describing in angular motion of SD2 relative to CD about S2s |
| $\theta_8$ | CD, SD2, SD1 | Term describing in angular motion of CD relative to SD2 about SD1 |
| Dihedral | | |
| $\Phi_1$ | SD2, SD1, RBD, RBDa | Dihedral describing rotation of SD2 and RBDa about SD1 to RBD axis |
| $\Phi_2$ | NTDa, NTD, NTD′, SD2 | Dihedral describing rotation of NTDa and SD2 about NTD to NTD′ axis |
| $\Phi_3$ | NTD′, SD2, SD1, RBD | Dihedral describing rotation of NTD′ and RBD about SD2 to SD1 axis |
| $\Phi_4$ | NTD′, SD2, CD, S2s | Dihedral describing rotation of NTD′ and S2s about SD2 to CD axis |
| $\Phi_5$ | NTD, NTD′, SD2, SD1 | Dihedral describing rotation of NTD and SD1 about SD2 to NTD′ axis |
| $\Phi_6$ | NTD, NTD′, SD2, CD | Dihedral describing rotation of NTD and CD about SD2 to NTD′ axis |
| $\Phi_7$ | SD1, SD2, CD, S2s | Dihedral describing rotation of SD1 and S2s about SD2 to CD axis |

**Supplementary Table 8.** Inter-protomer vector descriptions

| Distance | Centroids | Description |
|---|---|---|
| $d_1'$ | $RBD_A$, $RBD_C$ | Term describing the distance between RBD from protomer A and C |
| $d_2'$ | $RBD_C$, $RBD_B$ | Term describing the distance between RBD from protomer B and C |
| $d_3'$ | $RBD_A$, $RBD_B$ | Term describing the distance between RBD from protomer A and C |
| $d_4'$ | $RBD_A$, $NTD_C$ | Term describing the distance between RBD from protomer A and NTD from protomer C |
| $d_5'$ | $RBD_C$, $NTD_B$ | Term describing the distance between RBD from protomer C and NTD from protomer B |
| $d_6'$ | $RBD_B$, $NTD_A$ | Term describing the distance between RBD from protomer B and NTD from protomer A |
| $d_7'$ | $SD1_A$, $NTD'_C$ | Term describing the distance between SD1 from protomer A and NTD′ from protomer C |
| $d_8'$ | $SD1_C$, $NTD'_B$ | Term describing the distance between SD1 from protomer C and NTD′ from protomer B |
| $d_9'$ | $SD1_B$, $NTD'_A$ | Term describing the distance between SD1 from protomer B and NTD′ from protomer A |
| Angle | | |
| $\theta_1'$ | $RBD_A$, $RBD_C$ ,$RBD_B$ | Term describing in angular motion of $RBD_A$ relative to $RBD_B$ about $RBD_C$ |
| $\theta_2'$ | $RBD_C$, $RBD_B$ ,$RBD_A$ | Term describing in angular motion of $RBD_C$ relative to $RBD_A$ about $RBD_B$ |
| $\theta_3'$ | $RBD_B$, $RBD_A$, $RBD_C$ | Term describing in angular motion of $RBD_B$ relative to $RBD_C$ about $RBD_A$ |
| Dihedral | | |
| $\Phi_1'$ | $RBDa_A$, $RBD_A$, $RBD_C$, $RBDa_C$ | Dihedral describing rotation of $RBDa_A$ and $RBDa_C$ about $RBD_A$ to $RBD_C$ axis |
| $\Phi_2'$ | $RBDa_C$, $RBD_C$, $RBD_B$, $RBDa_B$ | Dihedral describing rotation of $RBDa_C$ and $RBDa_C$ about $RBD_C$ to $RBD_B$ axis |
| $\Phi_3'$ | $RBDa_B$, $RBD_B$, $RBD_A$, $RBDa_A$ | Dihedral describing rotation of $RBDa_B$ and $RBDa_A$ about $RBD_B$ to $RBD_A$ axis |
| $\Phi_4'$ | $SD1a_A$, $SD1_A$, $NTD'_C$, $NTD'a_C$ | Dihedral describing rotation of $SD1a_A$ and $NTDa_C$ about $SD1_A$ to $NTD_C$ axis |
| $\Phi_5'$ | $SD1a_C$, $SD1_C$, $NTD'_B$, $NTD'a_B$ | Dihedral describing rotation of $SD1a_C$ and $NTD'a_B$ about $SD1_C$ to $NTD'_B$ axis |
| $\Phi_6'$ | $SD1a_B$, $SD1_B$, $NTD'_A$, $NTD'a_A$ | Dihedral describing rotation of $SD1a_B$ and $NTD'a_A$ about $SD1_B$ to $NTD'_A$ axis |
| $\Phi_7'$ | $SD2a_A$, $SD1_A$, $NTD'_C$, $SD2a_C$ | Dihedral describing rotation of $SD2a_A$ and $SD2a_C$ about $SD1_A$ to $NTD'_C$ axis |
| $\Phi_8'$ | $SD2a_C$, $SD1_C$, $NTD'_B$, $SD2a_B$ | Dihedral describing rotation of $SD2a_C$ and $SD2a_B$ about $SD1_C$ to $NTD'_B$ axis |
| $\Phi_9'$ | $SD2a_B$, $SD1_B$, $NTD'_A$, $SD2a_A$ | Dihedral describing rotation of $SD2a_B$ and $SD2a_A$ about $SD1_B$ to $NTD'_A$ axis |

**Supplementary Table 9**. Statistics for 3D reconstruction and refinement for BA.3.2.1 spike local refinement model

| | $RBD_C$ /$RBD_A$/$NTD_B$ Closed spike | $RBD_A$/$NTD_C$ Closed spike | $RBD_B$/$NTD_A$ Closed spike | $NTD_B$ Closed spike | RBD/hACE2 Conformation 1 |
|---|---|---|---|---|---|
| EMD | 73599 | 73597 | 73619 | 73795 | 73426 |
| PDB | 9YX8 | 9YX7 | 9YXX | 9Z3Y | 9YSR |
| **Refinement** | | | | | |
| Symmetry imposed | C1 | C1 | C1 | C1 | C1 |
| Particles (no.) | 673,009 | 673,009 | 673,009 | 673,009 | 278,738 |
| Map resolution (Å) | 2.9 | 2.9 | 3.0 | 3.0 | 3.6 |
| **Model composition** | | | | | |
| Chains | 3 | 2 | 2 | 1 | 2 |
| Atoms | 4,848 | 3,609 | 3,737 | 1,936 | 6,273 |
| Residues (Protein) | 585 | 434 | 453 | 255 | 757 |
| Ligands (NAG) | 6 | 6 | 6 | 7 | 6 |
| **R.m.s. deviations** | | | | | |
| Bond lengths (Å) | 0.004 | 0.003 | 0.004 | 0.003 | 0.003 |
| Bond angles (°) | 0.622 | 0.613 | 0.623 | 0.633 | 0.654 |
| **Model statistics** | | | | | |
| Clash score | 6.65 | 4.23 | 4.90 | 5.77 | 9.14 |
| MolProbity score | 1.63 | 1.53 | 1.69 | 1.69 | 1.71 |
| Rotamer outliers (%) | 0.76 | 0.51 | 0.74 | 0.49 | 0.60 |
| **Ramachandran plot** | | | | | |
| Outliers (%) | 0.00 | 0.00 | 0.00 | 0.00 | 0.00 |
| Allowed (%) | 3.85 | 4.74 | 6.58 | 5.48 | 3.47 |

**Supplementary Table 10.** Statistics for 3D reconstruction and model refinement for LP.8.1 local refinement.

| | $RBD_A/NTD_C$ closed spike | $RBD_B/NTD_A$ closed spike | $RBD_C/NTD_B$ closed spike | RBD/hACE2 complex conformation 1 |
|---|---|---|---|---|
| EMD | 73441 | 73443 | 73444 | 73429 |
| PDB | 9YT4 | 9YT6 | 9YT7 | 9YSS |
| **Refinement** | | | | |
| Symmetry imposed | C1 | C1 | C1 | C1 |
| Particles (no.) | 948,866 | 948,866 | 948,866 | 232,292 |
| Map resolution (Å) | 2.8 | 2.8 | 2.8 | 3.7 |
| **Composition** | | | | |
| Chains | 2 | 2 | 2 | 2 |
| Atoms | 4,229 | 4,192 | 4,205 | 6,312 |
| Residues (Protein) | 510 | 508 | 510 | 761 |
| Ligands (NAG) | 11 | 9 | 9 | 8 |
| **R.m.s. deviations** | | | | |
| Bond lengths (Å) | 0.004 | 0.003 | 0.004 | 0.004 |
| Bond angles (°) | 0.654 | 0.582 | 0.690 | 0.717 |
| **Model statistics** | | | | |
| Clash score | 3.97 | 5.83 | 6.05 | 8.60 |
| MolProbity score | 1.36 | 1.50 | 1.45 | 1.69 |
| Rotamer outliers (%) | 0.22 | 0.22 | 0.22 | 0.75 |
| **Ramachandran plot** | | | | |
| Outliers (%) | 0.00 | 0.00 | 0.00 | 0.00 |
| Allowed (%) | 3.02 | 3.04 | 2.62 | 3.44 |
| Favored (%) | 97.18 | 96.96 | 97.38 | 96.56 |
